# Photosynthetic sea slugs inflict protective changes to the light reactions of the chloroplasts they steal from algae

**DOI:** 10.1101/2020.04.09.034124

**Authors:** Vesa Havurinne, Esa Tyystjärvi

**Author notes:** Corresponding author: Esa Tyystjärvi.

## Abstract

Sacoglossan sea slugs are able to maintain functional chloroplasts inside their own cells, and mechanisms that allow preservation of the chloroplasts are unknown. We found that the slug *Elysia timida* inflicts changes to the photosynthetic light reactions of the chloroplasts it steals from the alga *Acetabularia acetabulum*. Working with a large continuous laboratory culture of both the slugs (>500 individuals) and their prey algae, we show that the plastoquinone pool of slug chloroplasts remains oxidized, which can suppress reactive oxygen species formation. Slug chloroplasts also rapidly build up a strong proton motive force upon a dark-to-light transition, which helps them to rapidly switch on photoprotective non-photochemical quenching of excitation energy. Finally, our results suggest that chloroplasts inside *E. timida* rely on flavodiiron proteins as electron sinks during rapid changes in light intensity. These photoprotective mechanisms are expected to contribute to the long-term functionality of the chloroplasts inside the slugs.

## Introduction

The sea slug *Elysia timida* is capable of stealing chloroplasts from their algal prey (Figure 1). Once stolen, the chloroplasts, now termed kleptoplasts, remain functional inside the slug’s cells for several weeks, essentially creating a photosynthetic slug. The only animals capable of this phenomenon are sea slugs belonging to the Sacoglossan clade (Rumpho et al., 2011; de Vries et al., 2014). Despite decades of research, there is still no consensus about the molecular mechanisms that allow the slugs to discriminate other cellular components of the algae and only incorporate the chloroplasts inside their own cells, or how the slugs maintain the chloroplasts functional for times that defy current paradigms of photosynthesis. Also the question whether the slugs in fact get a real nutritional benefit from the photosynthates produced by the stolen chloroplasts, is still being debated (Cartaxana et al., 2017; Rauch et al., 2017).

**Figure 1.**
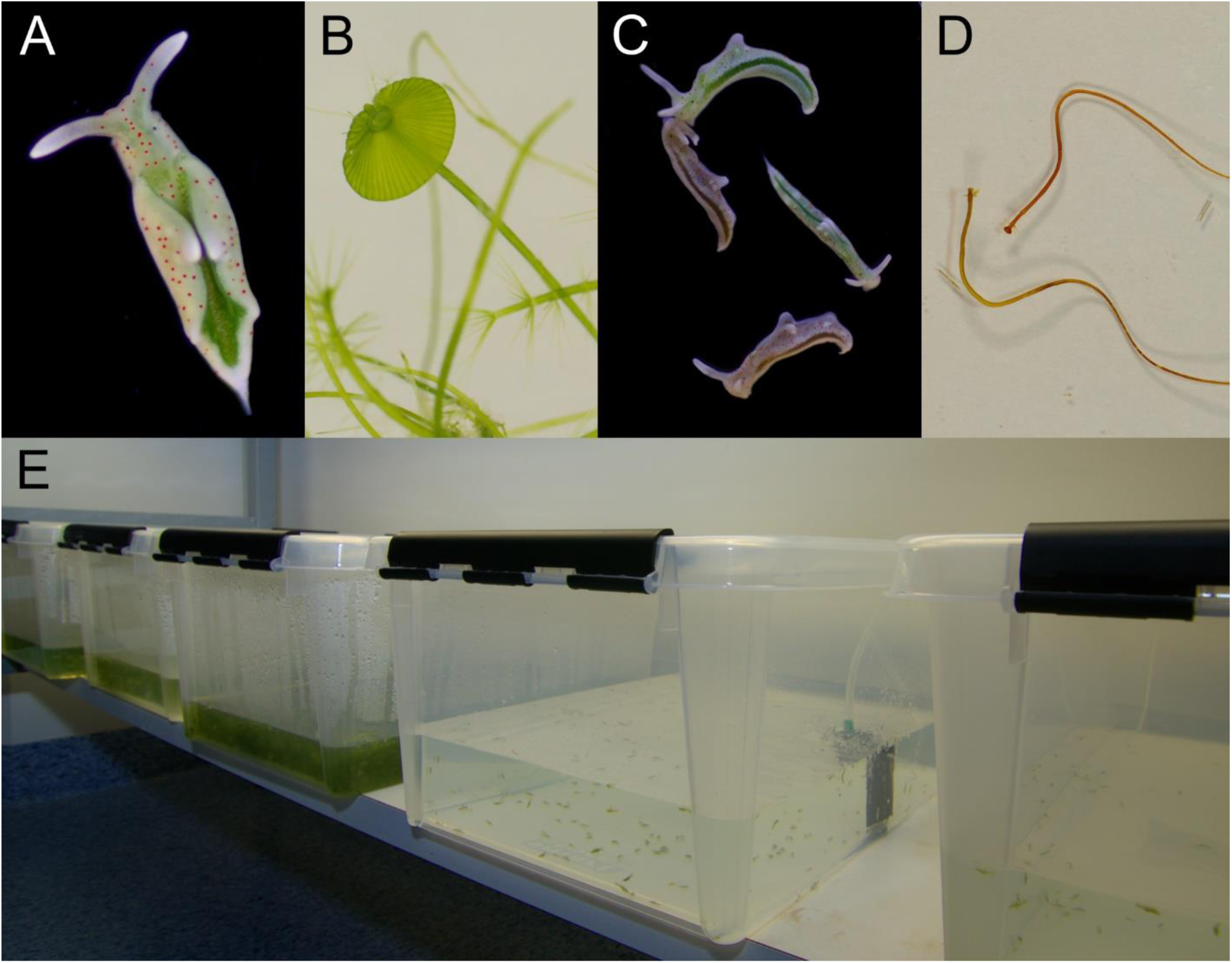
Laboratory cultures of the photosynthetic sea slug *E. timida* and its prey alga *Acetabularia*. A) A freshly fed adult *E. timida* individual. B) The giant-celled green alga *Acetabularia*. The cap-like structures are the site of gamete maturation and serve as indicators of the end of the vegetative growth phase of individual *Acetabularia* cells. C) The red morphotype *E. timida* can be induced by feeding it with red morphotype *Acetabularia*. Green morphotype *E. timida* individuals are also shown for reference. D) Red morphotype *Acetabularia* can be induced by subjecting the cells to cold temperature and high light (see “Materials and Methods” for details). E) *E. timida* and *Acetabularia* can be cultured in transparent plastic tanks. The two tanks in the foreground are *E. timida* tanks and the three other tanks contain *Acetabularia* cultures.

One of the main problems that kleptoplasts face is light induced damage to both photosystems. Photoinhibition of Photosystem II (PSII) takes place at all light intensities and photosynthetic organisms have developed an efficient PSII repair cycle to counteract it (Tyystjärvi, 2013). Unlike higher plants (Järvi et al., 2015), the chloroplast genomes of all algal species involved in long term kleptoplasty encode FtsH, a protease involved in PSII repair cycle (de Vries et al., 2013). However, out of all prey algae species of photosynthetic sea slugs, only in *Vaucheria litorea*, the prey alga of *Elysia chlorotica*, the chloroplast-encoded FtsH contains the critical M41 metalloprotease domain required for degradation of the D1 protein during PSII repair (Christa et al., 2018). Photoinhibition of Photosystem I (PSI) occurs especially during rapid changes in light intensity (Tikkanen and Grebe, 2018) and should cause problems in isolated chloroplasts in the long run. In addition to the specific inhibition mechanisms of the photosystems, unspecific damage caused by reactive oxygen species (ROS) (Khorobrykh et al., 2020) is expected to deteriorate an isolated chloroplast.

Photoprotective mechanisms counteract photodamage. Recent efforts have advanced our understanding of photoprotection in kleptoplasts (Christa et al., 2018; Cartaxana et al., 2019). It has been shown that kleptoplasts of *E. timida* do retain the capacity to induce physiological photoprotection mechanisms similar to the ones in the prey green alga *Acetabularia acetabulum* (hereafter *Acetabularia*) (Christa et al., 2018). The most studied mechanism is the “energy-dependent” qE component of non-photochemical quenching of excitation energy (NPQ). qE is triggered through acidification of the thylakoid lumen by protons pumped by the photosynthetic electron transfer chain. The xanthophyll cycle enhances qE (Papageorgiou and Govindjee, 2014) but there have been contrasting reports on the capability of *E. timida* to maintain a highly functional xanthophyll cycle if the slugs are not fed with fresh algae (i.e. starved) and about the effect of NPQ on kleptoplast longevity (Christa et al., 2018; Cartaxana et al., 2019). Although advancing our understanding of the mechanisms of kleptoplast longevity, these recent publications underline the trend of contradictory results that has been going on for a long time.

There are reports of continuous husbandry of photosynthetic sea slugs, mainly *E. timida* (Schmitt et al., 2014) and *E. chlorotica* (Rumpho et al., 2011), but still today most research is conducted on animals caught from the wild. We have grown the sea slug *E. timida* and its prey *Acetabularia* in our lab for several years (Figure 1). As suggested by Schmitt et al. (2014), *E. timida* is an attractive model organism for photosynthetic sea slugs because it is easy to culture with relatively low costs (Figure 1E). A constant supply of slugs has opened a plethora of experimental setups yet to be tested, one of the more exciting ones being the case of red morphotypes of both *E. timida* and *Acetabularia* (Figure 1C,D). Red morphotypes of *E. timida* and *Acetabularia* were first described by González-Wangüemert et al. (2006) and later shown to be due to accumulation of an unidentified carotenoid during cold/high-light acclimation of the algae that were then eaten by *E. timida* (Costa et al., 2012). The red morphotypes provide a visual proof that the characteristics of the kleptoplasts inside *E. timida* can be modified by acclimating their feedstock to different environmental conditions.

We optimized a completely new set of biophysical methods to study photosynthesis in the sea slugs and found differences in photosynthetic electron transfer reactions between *E. timida* and *Acetabularia* grown in varying culture conditions. The most dramatic differences between the slugs and their prey were noticed in PSII electron transfer of the red morphotype *E. timida* (Figure 1C) and *Acetabularia* (Figure 1D). In addition to measuring chlorophyll *a* fluorescence decay kinetics, we also measured fluorescence induction kinetics, PSI electron transfer and formation of proton motive force during dark to light transition. Our results suggest that dark reduction of the plastoquinone (PQ) pool, a reserve of central electron carriers of the photosynthetic electron transfer chain, is weak in the slugs compared to the algae, and that a strong build-up of proton motive force is likely linked to fast induction and elevated levels of NPQ in kleptoplasts. It is also clear that PSI utilizes oxygen sensitive flavodiiron proteins (FLVs) as alternative electron sinks in both the slugs and the algae, and this sink protects the photosynthetic apparatus from light-induced damage in *E. timida*.

## Results

### Non-photochemical reduction of the PQ pool is inefficient in *E. timida*

We estimated reoxidation kinetics of the first stable electron acceptor of PSII, Q_A_^-^, from dark acclimated *E. timida* and *Acetabularia* by measuring the decay of chlorophyll *a* fluorescence yield after a single turnover flash (Figure 2). In the green morphotypes of the slugs and the algae, Q_A_^-^ reoxidation kinetics were similar between the two species in aerobic conditions both in the absence and presence of 3-(3, 4-dichlorophenyl)-1, 1-dimethylurea (DCMU), an inhibitor of PSII electron transfer (Figure 2A,B). This indicates that electron transfer within PSII functions in the same way in both species. In anaerobic conditions, fluorescence decay was slower than in aerobic conditions in both species (Figure 2C), suggesting that the environment of the slug kleptoplasts normally remains aerobic in the dark even in the presence of slug respiration. Decay of fluorescence is slow in anaerobic conditions probably because reduced PQ, accumulating in the dark, hinders Q_A_^-^ reoxidation (de Wijn and van Gorkom, 2001; Oja et al., 2011; Deák et al., 2014; Krishna et al., 2019). Anaerobicity slowed the fluorescence decay less in *E. timida* than in *Acetabularia*, especially during the fast (∼ 300–500 μs) and middle (∼ 5–15 ms) phases of fluorescence decay in anaerobic conditions (Figure 2C). This could indicate that non-photochemical reduction of the PQ pool during the dark acclimation period is less efficient in the slug than in the alga.

**Figure 2.**
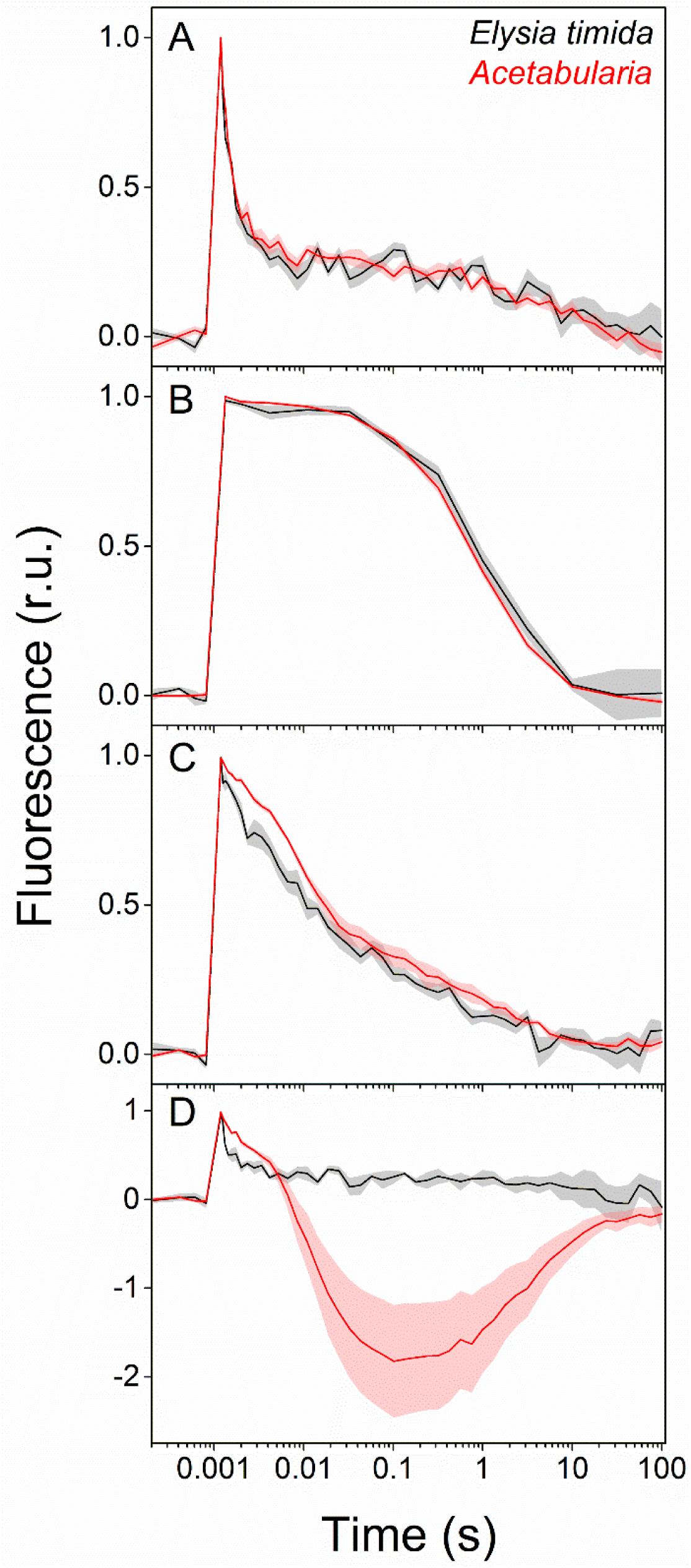
Differences in the redox poise of the PQ pool after dark acclimation lead to differences in Q_A_^-^ reoxidation between *E. timida* (black) and *Acetabularia* (red). A-C) Chlorophyll *a* fluorescence yield decay after a single turnover flash in aerobic conditions without any inhibitors in regular, green morphotypes of *E. timida* and *Acetabularia* (A), in aerobic conditions in the presence of 10 µM DCMU (B), and in anaerobic conditions, achieved by a combination of glucose oxidase (8 units/ml), glucose (6 mM) and catalase (800 units/ml), in the absence of inhibitors (C). See Figure 2 – figure supplement 1A for details on the anaerobic conditions. D) Chlorophyll fluorescence decay measured from the red morphotypes of *E. timida* and *Acetabularia* in aerobic conditions without any inhibitors. Fluorescence traces were double normalized to their respective minimum (measured prior to the single turnover flash), and maximum fluorescence levels. Curves in (A) are averages from 7 (*E. timida*) and 5 (*Acetabularia*) biological replicates, 4 and 5 in (B), and 5 and 5 in (C-D), respectively. The shaded areas around the curves represent SE. All *E. timida* data are from individuals taken straight from the feeding tanks without an overnight starvation period. See Figure 2 – source data 1 for original data.

Differences in dark reduction of the PQ pool between the slugs and the algae were also supported by Q_A_^-^ reoxidation measurements from the red morphotype *E. timida* and *Acetabularia* (see Figure 1C,D for images of the red morphotypes and “Materials and methods” for their preparation). Fluorescence decay in red *Acetabularia* followed very strong wave like kinetics, with a large undershoot below the dark-acclimated minimum fluorescence level, while there was no sign of such kinetics in the red morphotype *E. timida* (Figure 2D). The wave phenomenon has been characterized in detail in several species of cyanobacteria, where anaerobic conditions in the dark are enough for its induction (Wang et al., 2012; Deák et al., 2014; Ermakova et al., 2016). According to Deák et al. (2014), anaerobic conditions cause a highly reduced PQ pool through respiratory electron donation, mainly from NAD(P)H that is mediated by the NAD(P)H-dehydrogenase (NDH). The wave phenomenon was recently also characterized in the green alga *Chlamydomonas reinhardtii* (Krishna et al., 2019). The authors suggested that, similar to cyanobacteria, non-photochemical reduction of the PQ pool by stromal reductants in anaerobic conditions in the dark leads to wave like kinetics of fluorescence decay after a single turnover flash in sulphur deprived *C. reinhardtii* cells. The exact mechanisms underlying the wave phenomenon are still unknown, but the involvement of the NDH complex is clear (Deák et al., 2014; Krishna et al., 2019). Chemical inhibition of the NDH complex in *C. reinhardtii* led to complete abolishment of the wave phenomenon in cells that were otherwise primed for it (Krishna et al., 2019). Interestingly, the comparison of NDH uninhibited and inhibited cells resulted in fluorescence decay kinetics that are highly reminiscent of the kinetics in Figure 2D, with the red morphotype *E. timida* and red morphotype *Acetabularia* being analogous to the NDH inhibited and uninhibited *C. reinhardtii* cells, respectively.

### Full photochemical reduction of the PQ pool is delayed in *E. timida*

In order to investigate whether the alterations in the dark reduction of the PQ pool in kleptoplasts lead to differences in electron transfer reactions during continuous illumination, we measured chlorophyll *a* fluorescence induction kinetics from both *E. timida* and *Acetabularia* (Figure 3). Briefly, fluorescence rise during the first ∼1 s of continuous illumination of a photosynthetic sample can be divided into distinct phases, denoted as O-J-I-P, when plotted on a logarithmic time scale (see Figure 3A). Alterations in the magnitude and time requirements of these phases are indicative of changes in different parts of the photosynthetic electron transfer chain (Strasser et al., 1995; Strasser et al., 2004; Kalaji et al., 2014).

**Figure 3.**
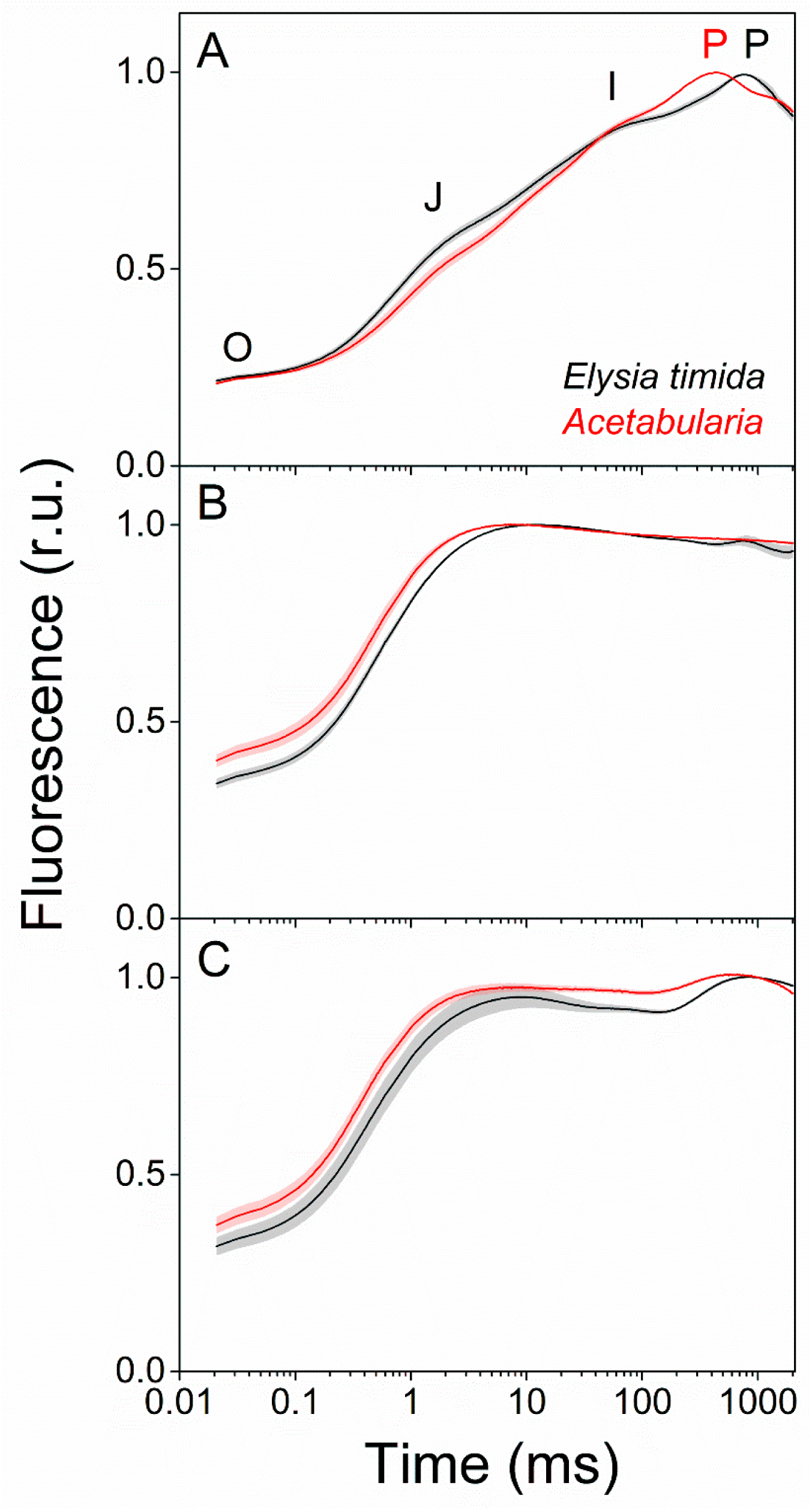
Fluorescence induction kinetics during dark-to-light transition indicate differences in full photochemical reduction of the PQ pool between *E. timida* (black) and *Acetabularia* (red). A-C) Multiphase chlorophyll *a* fluorescence induction transient (OJIP) measured from dark acclimated *E. timida* and *Acetabularia* in aerobic conditions without any inhibitors (A), in the presence of 10 µM DCMU (B), and in anaerobic conditions, achieved by a combination of glucose oxidase (8 units/ml), glucose (6 mM) and catalase (800 units/ml), without any inhibitors (C). Fluorescence traces were normalized to their respective maximum fluorescence levels. For unnormalized data, see Figure 3 – figure supplement 1B. Curves in (A) are averages from 10 (*E. timida*) and 12 (*Acetabularia*) biological replicates, 10 and 9 in (B), and 13 and 11 in (C), respectively. The shaded areas around the curves represent SE. All *E. timida* data are from individuals taken straight from the feeding tanks without an overnight starvation period. See Figure 3 – source data 1 for original data.

Fluorescence induction measurements in aerobic conditions revealed that in green *E. timida* individuals, maximum fluorescence (P phase of OJIP fluorescence rise kinetics) was reached ∼300 ms later than in green *Acetabularia* (Figure 3A). To investigate whether the elongated time requirement to reach maximum fluorescence is caused by light attenuation in the slug tissue, we tested the effect of different intensities of the light pulse to the fluorescence transient in *E. timida*. In the tested range, the intensity of the pulse did affect the O-J-I phases but not the time of the P phase in aerobic conditions (Figure 3 - figure supplement 1). This suggests that the ∼300 ms delay of the P phase in *E. timida* is of physiological origin. The delay in fluorescence induction in *E. timida* in Figure 3A could indicate that the PQ pool is more oxidized in the slugs than in *Acetabularia* even in aerobic conditions, and full reduction of the PQ pool simply takes longer in *E. timida*. We base this on the notion that full reduction of the PQ pool is a prerequisite for reaching maximum fluorescence when fluorescence rise kinetics are measured using multiple turnover saturating light pulses, such as the ones used in the current study (Kramer et al., 1995; Yaakoubd et al., 2002; Suggett et al., 2003; Osmond et al., 2017). It should be noted, however, that the mechanisms controlling the J-I-P phase (also known as the thermal phase) of fluorescence induction are still under debate (Stirbet and Govindjee, 2012; Schansker et al., 2014; Vredenberg, 2015; Havurinne et al., 2018; Magyar et al., 2018; Schreiber et al., 2019).

We witnessed a slightly slower O-J phase in DCMU treated *E. timida* individuals (∼10 ms to reach J) than in DCMU treated *Acetabularia* (∼6 ms) (Figure 3B). The O-J phase is considered to represent the reduction of Q_A_ to Q_A_^-^, and it is indicative of the amount of excitation energy reaching PSII, i.e. functional absorption cross-section of the PSII light harvesting antennae (Kalaji et al., 2014). The J-I-P phases are nullified when DCMU is introduced into the sample, as forward electron transfer from Q_A_^-^ is blocked (Kodru et al., 2015), which makes the O-J phase highly distinguishable. It is possible that the slower O-J fluorescence rise indicates a decrease in the functional absorption cross-section of PSII in the slug cells (Figure 3B). In anaerobic conditions the OJIP transient behaved in a manner that can be explained by blockages in the electron transfer chain, i.e. a highly reduced PQ pool (Figure 3C). This blockage seems to affect the electron transfer more in *Acetabularia* than in *E. timida*, i.e. the J-I-P phases are more pronounced in *E. timida*, supporting the earlier suggestion that in anaerobic conditions electron transfer from Q_A_^-^ to Q_B_ and PQ pool is faster in *E. timida* than in *Acetabularia*.

### Build-up of proton motive force in *E. timida* may facilitate rapid induction of NPQ

To inspect intricate differences in proton motive force formation between *E. timida* and *Acetabularia*, we measured electrochromic shift (ECS) from dark acclimated individuals of both species during a strong, continuous light pulse (Figure 4A). According to the ECS data, proton motive force of *Acetabularia* dissipates to a steady level after the initial spike in thylakoid membrane energization, whereas in *E. timida* there is a clear build-up of proton motive force after a slight relaxation following the initial spike. The build-up in *E. timida* suggests that protons are released into the thylakoid lumen during illumination, but not out. This could be indicative of defects in ATP-synthase functionality in the slugs, perhaps due to lack of inorganic phosphate. Furthermore, while the capacity to induce photoprotective NPQ in *E. timida* has been shown to reflect the acclimation state of its food source, *E. timida* and also *E. chlorotica* consistently exhibit higher levels of NPQ than their respective algal food sources (Cruz et al., 2015; Christa et al., 2018; also see below “P700 redox kinetics in *E. timida* are affected by the acclimation status of its prey”). This is likely linked to the stronger acidification of the lumen in the slugs (Figure 4A), since the major qE component of NPQ is pH dependent (Müller et al., 2001; Papageorgiou and Govindjee, 2014).

**Figure 4.**
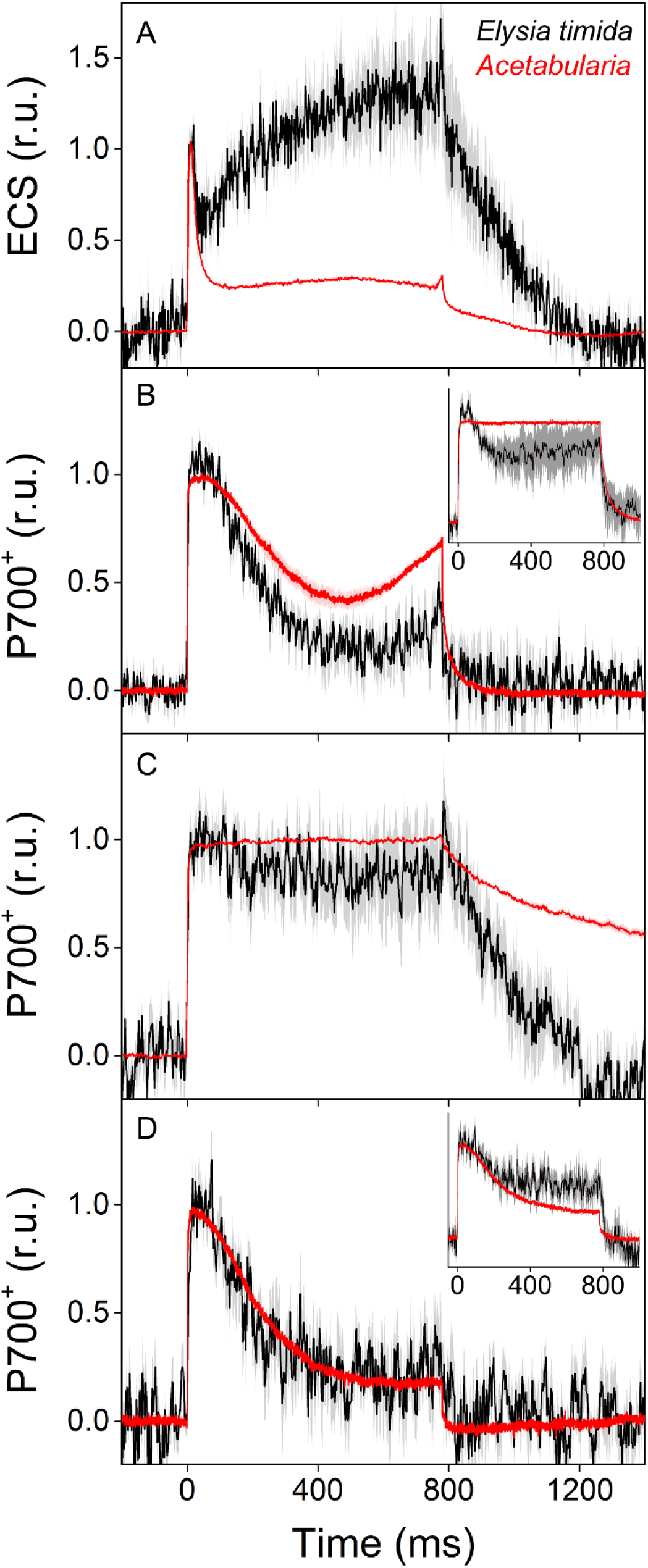
ECS and P700^+^ measurements indicate differences in proton motive force formation and utilization of alternative electron acceptors of PSI between dark acclimated *E. timida* (black) and *Acetabularia* (red) during a 780 ms high-light pulse. A) ECS measured from *E. timida* and *Acetabularia* upon exposure to a high-light pulse in aerobic conditions. B) P700 redox kinetics upon exposure to a high-light pulse in aerobic conditions without any inhibitors. The inset shows P700 redox kinetics from the same samples during a second high-light pulse, fired 10 s after the first one. C) P700 oxidation kinetics in the presence of 10 µM DCMU. D) P700 oxidation kinetics in the absence of DCMU in anaerobic conditions, achieved by a combination of glucose oxidase (8 units/ml), glucose (6 mM) and catalase (800 units/ml). The inset shows P700 oxidation kinetics from the same samples during the second high-light pulse. ECS and P700^+^ transients were double normalized to their respective dark levels (measured prior to the onset of the high-light pulse), and to the initial ECS or P700^+^ peak (measured immediately after the onset of the pulse). Curves in (A) are averages from 13 (*E. timida*) and 6 (*Acetabularia*) biological replicates, 7 and 3 in (B), 13 and 4 in (C), and 8 (7 in inset) and 3 in (D), respectively. The shaded areas around the curves represent SE. All *E. timida* data are from individuals taken straight from the feeding tanks, without an overnight starvation period. See Figure 4 – source data 1 for original data.

### Flavodiiron proteins function as alternative electron sinks in *E. timida* and *Acetabularia*

We utilized a nearly identical protocol as Shimakawa et al. (2019) to measure redox kinetics of P700, the reaction center chlorophyll of PSI, in dark acclimated *E. timida* and *Acetabularia* during dark-to-light transition (Figure 4B-D). In aerobic conditions, *Acetabularia* P700 redox kinetics during a high-light pulse followed the scheme where P700 is first strongly oxidized due to PSI electron donation to downstream electron acceptors such as ferredoxin, and re-reduced by electrons from the upstream electron transfer chain (Figure 4B). Finally, oxidation is resumed by alternative electron acceptors of PSI, most likely FLVs, as they have been shown to exist in all groups of photosynthetic organisms except angiosperms and certain species belonging to the red-algal lineage (Allahverdiyeva et al., 2015; Ilík et al., 2017; Shimakawa et al., 2019). P700 redox kinetics in *E. timida* and *Acetabularia* in aerobic conditions were similar in terms of the overall shape of the curve, but the re-oxidation phase after ∼600 ms was dampened in *E. timida* (Figure 4B). In the presence of DCMU, P700 remained oxidized throughout the pulse in both species (Figure 4C). This shows that the re-reduction of P700^+^ after the initial peak in aerobic conditions (Figure 4B) is due to electron donation from PSII in *E. timida* and *Acetabularia*. The complete absence of the final oxidation phase in both species in anaerobic conditions (Figure 4D) supports the view that FLVs are indeed behind P700 oxidation during the high-light pulse and function as electron sinks in both *E. timida* and *Acetabularia* by donating electrons to oxygen (Ilík et al., 2017; Shimakawa et al., 2019).

During the optimization process of the P700 oxidation measurements, we found that firing a second high-light pulse 10 s after the first pulse resulted in a higher capacity to maintain P700 oxidized in both *E. timida* and *Acetabularia* in aerobic conditions (Figure 4B, inset). This procedure will hereafter be referred to as “second pulse protocol”. In *E. timida* the oxidation capacity was moderate even with the second pulse, showing a high re-reduction of P700^+^ after the initial oxidation. In *Acetabularia* the second pulse rescued P700 oxidation capacity completely. Full activation of FLVs as electron acceptors of PSI takes ∼1s after a transition from dark to light and they subsequently remain a considerable electron sink during the time required for light activation of Calvin-Benson-Bassham cycle (Ilík et al., 2017; Gerotto et al., 2016; Bulychev et al., 2018). The mechanism of such fast regulation of FLV functionality is not known, but it has been suggested that conserved cysteine residues of FLVs could offer a means for redox regulation through conformational changes (Alboresi et al., 2019). It is unclear whether the witnessed increase in P700 oxidation capacity during the second pulse is due to such activation of the FLVs. However, when a second pulse was fired in anaerobic conditions, P700 oxidation capacity showed only weak signs of improvement in *E. timida* and *Acetabularia* (Figure 4D, inset). It is therefore likely that FLVs are also behind P700 oxidation during the second pulse and the data in Figure 4B inset further support the suggestion that both *E. timida* and *Acetabularia* do utilize FLVs as electron sinks, although there are differences in their functionality between the two.

### P700 redox kinetics in *E. timida* are affected by the acclimation status of its prey

We tested the sensitivity of *E. timida* P700^+^ measurements by inflicting changes to the P700 oxidation capacity of its prey, and then estimating whether the changes are present in the slugs after feeding them with differently treated *Acetabularia*. First, we grew *Acetabularia* in elevated (1 %) CO_2_ environment, as high CO_2_ induces downregulation of certain FLVs in cyanobacteria (Zhang et al., 2012; Santana-Sanchez et al., 2019). Next, we allowed *E. timida* to feed on high-CO_2_ *Acetabularia* for four days. We used *E. timida* individuals that had been pre-starved for four weeks to ensure that the slugs would only contain chloroplasts from high-CO_2_ *Acetabularia*. After feeding, the slugs were allowed to incorporate the chloroplasts into their own cells for an overnight dark period in the absence of *Acetabularia* prior to the measurements. A similar treatment was applied to slug individuals that were fed ambient-air grown *Acetabularia* (see “Materials and methods” for the differences in the feeding regimes). These slugs will hereafter be termed as high-CO_2_ and ambient-air *E. timida*, respectively.

Ambient-air *E. timida* exhibited stronger P700 oxidation during the initial dark-to-light transition than high-CO_2_ *E. timida* (Figure 5A). When the second pulse protocol was applied, both groups showed a clear increase in P700 oxidation capacity, but once again P700 oxidation was stronger in the ambient-air slugs (Figure 5C). The differences between ambient-air and high-CO_2_ *Acetabularia* showed the same trend (Figure 5B,D). The simplest explanation for the decrease in P700 oxidation capacity in high-CO_2_ *E. timida* and *Acetabularia* is downregulation of FLVs. It is, however, possible that the differences are due to other changes caused by the high-CO_2_ acclimation, as, to our knowledge, the CO_2_ response of FLVs in green algae has not been studied in detail. Based on the highly similar ratio of chlorophylls *a* and *b* between ambient-air (chlorophyll *a*/*b*=2.37, SE±0.03, n=5) and high-CO_2_ slugs (chlorophyll *a*/*b*=2.41, SE±0.06, n=6), it seems likely that the acclimation process did not drastically alter the stoichiometry of PSII and PSI, which could have affected P700 oxidation during a high-light pulse.

**Figure 5.**
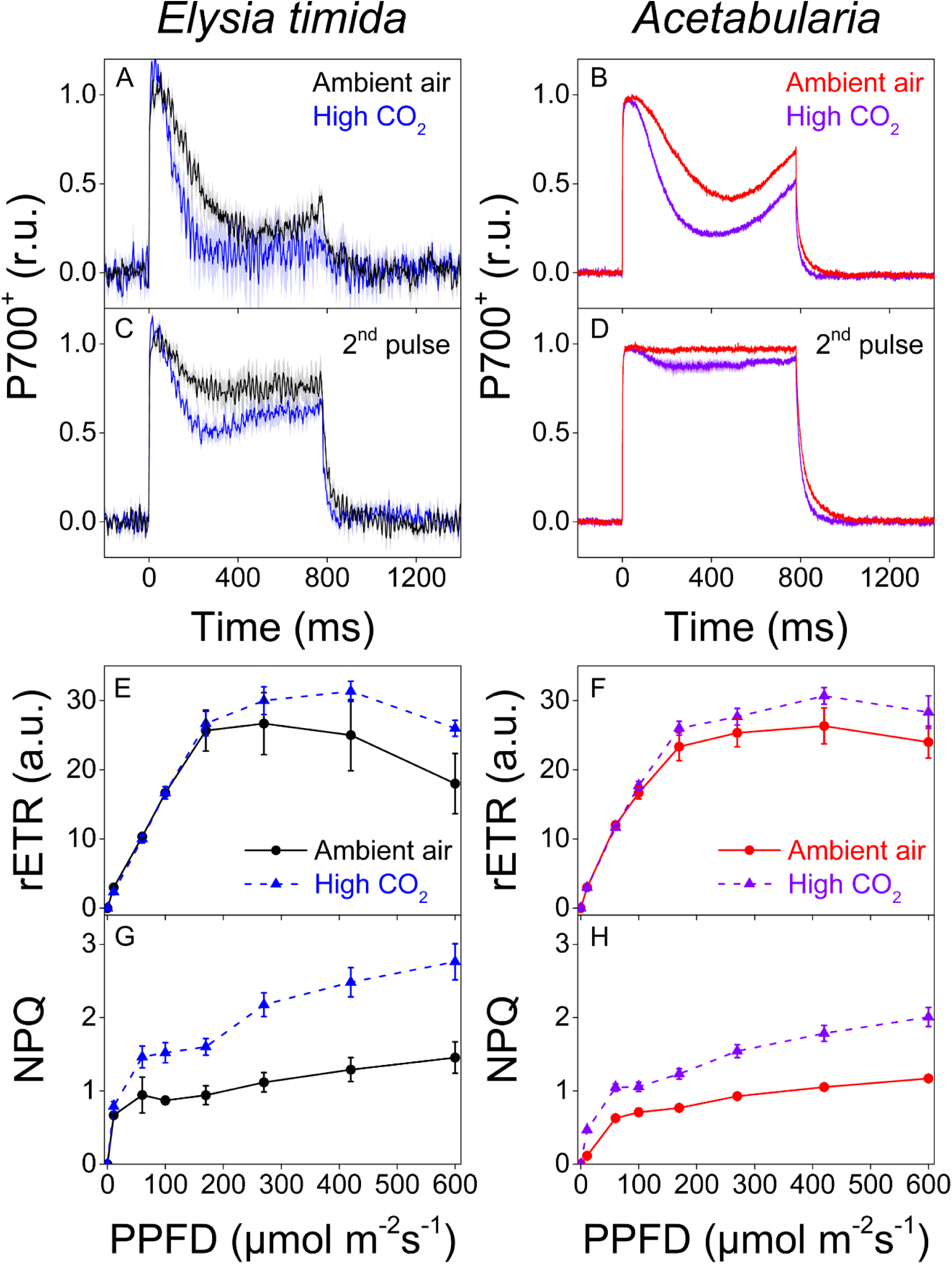
P700 redox kinetics, photosynthetic electron transfer and photoprotective NPQ induction in *E. timida* kleptoplasts (left panels) are affected by the CO_2_ acclimation state of its feedstock *Acetabularia* (right panels). A-B) P700 redox kinetics in dark acclimated ambient-air (black) and high-CO_2_ *E. timida* (blue) (A) and ambient-air (red) and high-CO_2_ *Acetabularia* (purple) (B) upon exposure to a 780 ms high-light pulse. The ambient-air *Acetabularia* data are the same as in Figure 4B and are shown here for reference. C-D) P700 redox kinetics during a second light pulse, fired 10 s after the first pulse, shown in panels A-B, in ambient-air and high-CO_2_ *E. timida* (C) and ambient-air and high-CO_2_ *Acetabularia* (D). The ambient-air *Acetabularia* data are the same as in Figure 4B inset and are shown here for reference. E-F) RLC measurements from dark acclimated ambient-air (black solid line) and high-CO_2_ *E. timida* (blue dashed line) (E) and ambient-air (red solid line) and high-CO_2_ *Acetabularia* (purple dashed line) (F). Illumination at each light intensity (PPFD) was continued for 90 s prior to firing a saturating pulse to determine relative electron transfer rate of PSII (rETR). *E. timida* individuals used in RLC measurements were fixed in 1 % alginate for the measurements (see “Materials and methods” and Figure 5 – figure supplement 1 for details). G-H) NPQ induction during the RLC measurements from ambient-air and high-CO_2_ *E. timida* (G) and ambient-air and high-CO_2_ *Acetabularia* (H). P700^+^ transients were double normalized to their respective dark levels and to the P700^+^ peak measured immediately after the onset of the pulse. Curves in (A) are averages from 7 (ambient-air *E. timida*) and 8 (high-CO_2_ *E. timida*) biological replicates, 7 and 8 in (C), and 3 and 3 in (E,G), respectively. High-CO_2_ *Acetabularia* curves in (B,D) are averages from 3 biological replicates. Ambient-air and high-CO_2_ *Acetabularia* curves in (F,H) are averages from 3 biological replicates. Shaded areas around the curves and error bars show SE. rETR and NPQ were calculated as described in “Materials and methods”. All *E. timida* individuals used in panels (A,C,E,G) were allowed to incorporate the chloroplasts for an overnight dark period in the absence of *Acetabularia* prior to the measurements. See Figure 5 – source data 1 for original data.

Acclimation to high CO_2_ also caused changes to PSII activity, estimated as relative electron transfer of PSII (rETR) during rapid light curve (RLC) measurements from dark acclimated samples (Figure 5E,F). Maximal rETR was lower in ambient-air *E. timida* and *Acetabularia* than in their high-CO_2_ counterparts. Also NPQ induction during the RLC measurements indicated that ambient-air and high-CO_2_ *E. timida* had very similar photosynthetic responses as their respective food sources (Figure 5G,H). However, the slugs exhibited faster induction and higher levels of NPQ than the algae, which could be due to stronger build-up of proton motive force in the kleptoplasts (see above “Build-up of proton motive force in E. timida may facilitate rapid induction of NPQ”). Because rETR in high light intensities (photosynthetic photon flux density, PPFD>200 µmol m^-2^s^-1^) was lower in ambient-air *E. timida* and *Acetabularia* (Figure 5E,F), it is possible that the stronger re-reduction of P700^+^ in high-CO_2_ *E. timida* and *Acetabularia* is due to more efficient electron donation to PSI. Whatever the exact reason behind the altered P700 oxidation kinetics is, it is clear that acclimation of *Acetabularia* to high CO_2_ lowers P700 oxidation capacity and this acclimation state is transferred into *E. timida*.

### High P700 oxidation capacity improves kleptoplast longevity under fluctuating light

Recently, there has been an increasing interest in protection of PSI by FLVs (Shimakawa et al., 2019; Gerotto et al., 2016; Jokel et al., 2018). This led us to investigate whether P700 oxidation capacity would affect the longevity of the kleptoplasts. We first compared kleptoplast longevity in ambient-air and high-CO_2_ *E. timida* in starvation under normal day/night cycle (12/12h, PPFD 40 µmol m^-2^s^-1^ during daylight hours), i.e. steady-light conditions. The slugs used here were subjected to a 4 week pre-starvation protocol prior to feeding them with their respective algae before the onset of the actual steady-light starvation experiment (see “P700 redox kinetics in *E. timida* are affected by the acclimation status of its prey” and “Materials and methods” for details).

Both groups behaved very similarly in terms of slug coloration and size, maximum quantum yield of PSII photochemistry (F_V_/F_M_) and minimal and maximal chlorophyll fluorescence (F_0_ and F_M_, respectively) during a 46 day starvation period (Figure 6). In both groups, F_V_/F_M_ decreased during starvation in a bi-phasic pattern, with slow decrease until day 21, after which PSII activity declined rapidly. The overall decline in F_V_/F_M_ was nearly identical in both groups throughout the experiment (Figure 6C). The initial population size of both groups was 50 slugs, and starvation induced deaths of 9 and 12 slugs during the experiment from ambient-air and high-CO_2_ *E. timida* populations, respectively (see “Materials and methods” for details on mortality and sampling). P700^+^ measurements from slugs starved for 5 days indicated that the starved ambient-air *E. timida* retained a higher P700 oxidation capacity through starvation than high-CO_2_ *E. timida*, when the second pulse P700^+^ kinetics protocol was applied (Figure 6E). These results show that altered P700 oxidation capacity does not affect chloroplast longevity in *E. timida* in steady-light conditions.

**Figure 6.**
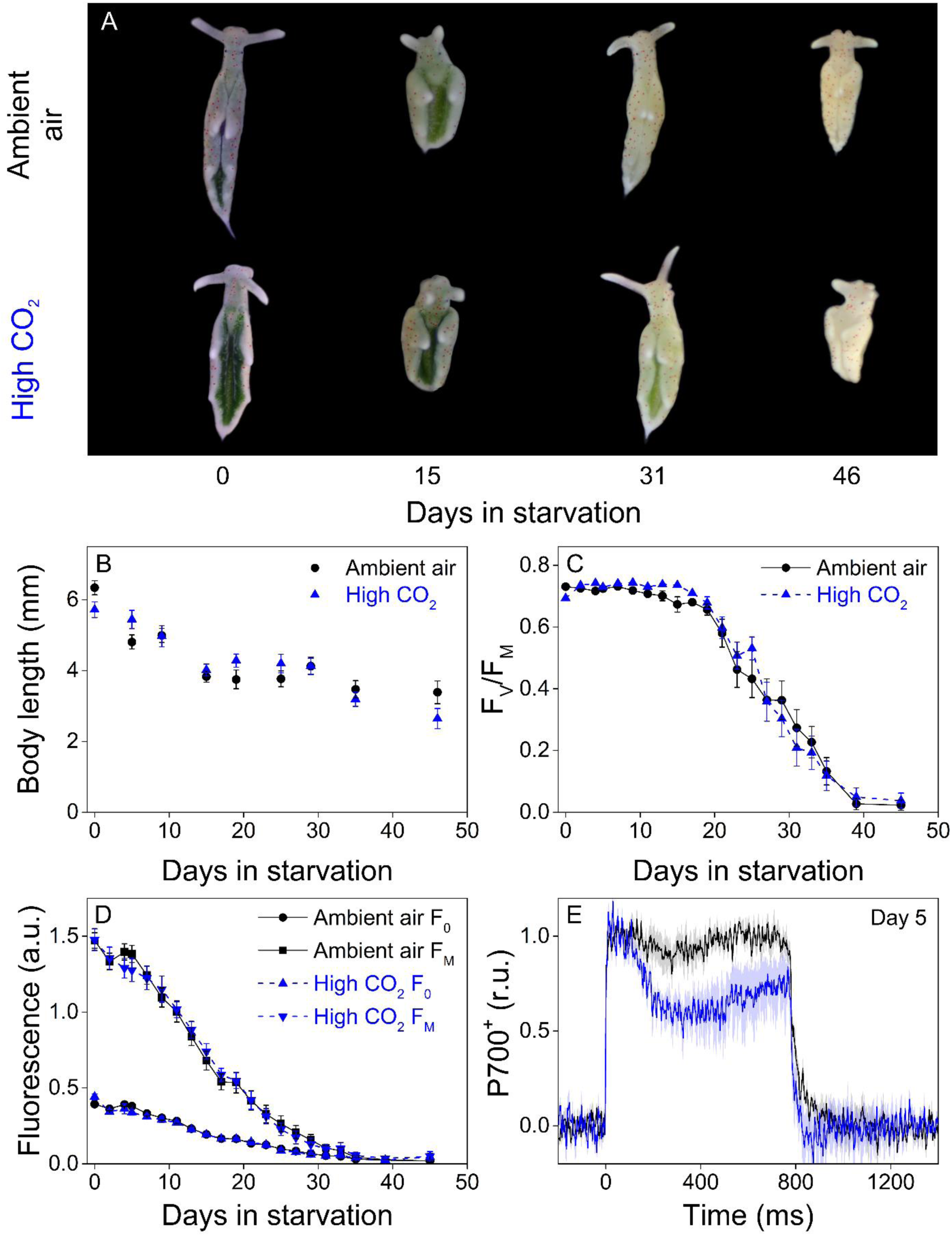
Altered P700 oxidation capacity does not affect chloroplast longevity in *E. timida* during starvation in steady-light conditions. A-B) Coloration of selected individuals (A) and body length (B) of the ambient-air (black) and high-CO_2_ *E. timida* (blue) slugs during steady-light starvation. The slug individuals in (A) do not show the actual scale of the slugs with respect to each other. C-D) Maximum quantum yield of PSII photochemistry (F_V_/F_M_) (C) and minimum (F_0_) and maximum chlorophyll *a* fluorescence (F_M_) (D) during starvation in ambient-air (black) and high-CO_2_ *E. timida* (blue). E) Second pulse P700 oxidation kinetics after five days in steady-light starvation in ambient-air (black) and high-CO_2_ *E. timida* (blue). Steady-light starvation light regime was 12/12h day/night and PPFD was 40 µmol m^-2^s^-1^ during daylight hours. See Figure 6 – figure supplement 1B for the spectra of lamps used in starvation experiments. All data in (B-D) represent averages from 50 to 8 biological replicates (see “Materials and methods” for details on mortality and sampling) and error bars show SE. P700^+^ transients in were double normalized to their respective dark levels and to the P700^+^ peak measured immediately after the onset of the pulse, and the curves in (E) represent averages from 7 (ambient air *E. timida*) and 5 biological replicates (high-CO_2_ *E. timida*) and the shaded areas around the curves show SE. See Figure 6 – source data 1 for original data from panels B-E.

We repeated the starvation experiment with new populations of ambient-air and high-CO_2_ *E. timida*, but this time the moderate background illumination (PPFD 40 µmol m^-2^s^-1^) was supplemented every 10 min with a 10 s high-light pulse (PPFD 1500 µmol m^-2^s^-1^) during daylight hours, i.e. fluctuating light. Slugs used in this experiment were not subjected to a pre-starvation protocol prior to feeding them with their respective algae but were simply allowed to replace their old chloroplasts with new specific ones during 6 days of feeding. The starting population size was 45 slugs for both groups and there were no starvation-caused losses during the whole experiment (see “Materials and methods” for details on sampling).

Starvation in fluctuating light induced faster onset of the rapid phase of F_V_/F_M_ decrease in both groups (Figure 7A) when compared to the steady light starvation experiment (Figure 6C). The exact onset of the rapid decline of F_V_/F_M_ was difficult to distinguish, but in ambient-air *E. timida* F_V_/F_M_ decrease accelerated only after ∼20 days, whereas in high-CO_2_ *E. timida* the turning point was during days 10-14 (Figure 7A). The overall longevity of PSII photochemistry in both groups was, however, very similar, as there was a sudden drop in F_V_/F_M_ in the ambient-air slugs on day 31. F_0_ behaved identically in the two groups, apart from days 0-4, when F_0_ of the high-CO_2_ *E. timida* dropped to the level of the ambient-air *E. timida* F_0_. At day 10, F_M_ of the high-CO_2_ slugs dropped drastically and the groups started to differ. Using the second pulse P700^+^ measurement protocol, we confirmed that the differences in P700 oxidation capacity were noticeable on days 0 and 6 in starvation also in the populations of slugs used for the fluctuating-light experiment (Figure 7C,D). RLC measurements were performed on the slugs on day 10 to inspect the underlying causes of the suddenly decreasing fluorescence parameters F_V_/F_M_ and F_M_ of the high-CO_2_ slugs (Figure 7E). The situation after 10 days in fluctuating light seemed almost the opposite to the situation on day 0 (Figure 5E), i.e. now the ambient-air slugs showed higher rETR_MAX_ than the high-CO_2_ slugs (Figure 7E). However, NPQ induction during the RLC measurements showed that high-CO_2_ *E. timida* were still able to generate and maintain stronger NPQ than ambient-air *E. timida* (Figure 7F), although the differences were not as strong as on day 0 (Figure 5G). These data indicate that the initial chloroplast acclimation status is retained during starvation, and the decrease in rETR_MAX_ in high-CO_2_ *E. timida* is likely due to light induced damage to the photosynthetic apparatus.

**Figure 7.**
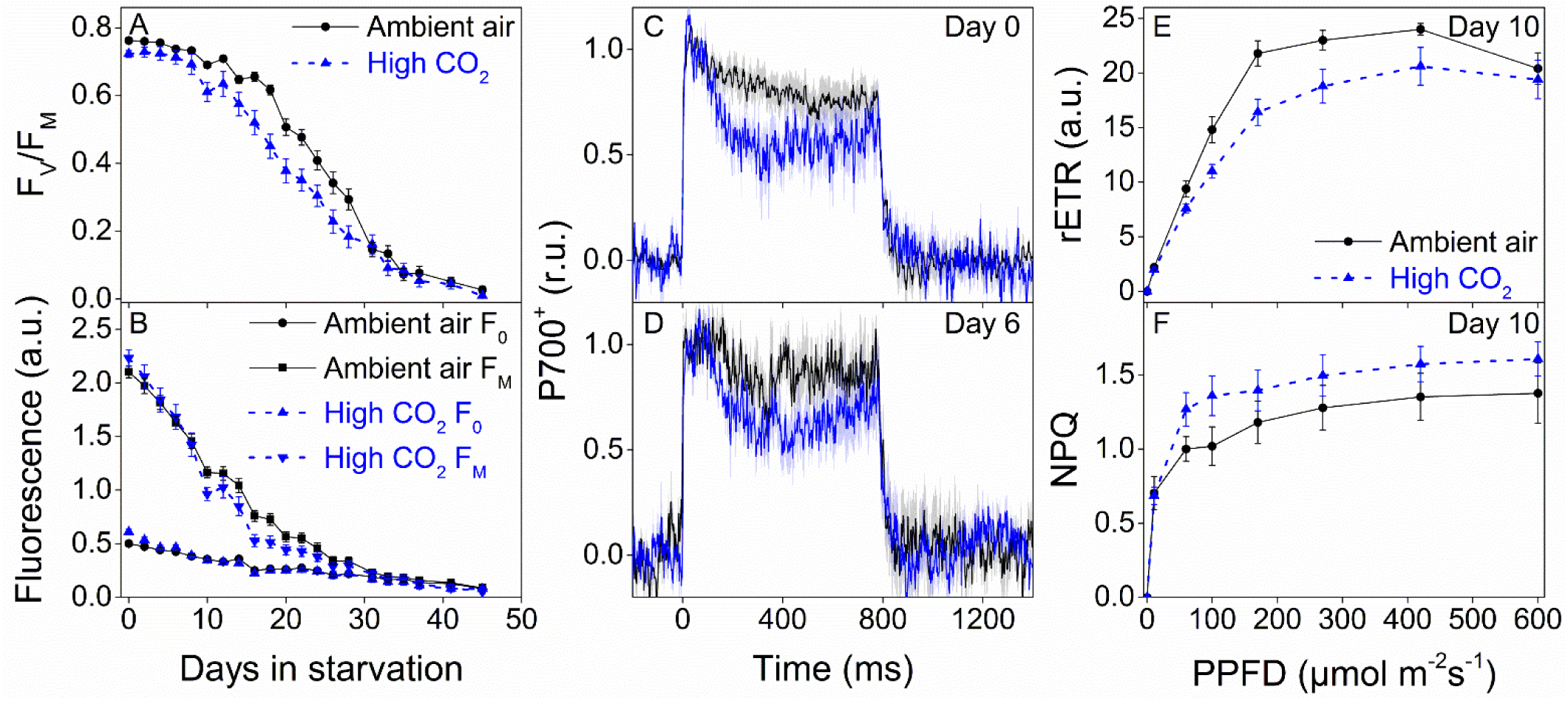
Higher P700 oxidation capacity protects the photosynthetic apparatus of ambient-air *E. timida* during fluctuating-light starvation. A-B) Maximum quantum yield of PSII photochemistry (F_V_/F_M_) (A) and minimum (F_0_) and maximum chlorophyll fluorescence (F_M_) (B) during fluctuating light starvation in ambient-air (black solid lines) and high-CO_2_ *E. timida* (blue dashed lines). C-D) Second pulse P700 oxidation kinetics after 0 and 5 days in fluctuating-light starvation in ambient-air (black) and high-CO_2_ *E. timida* (blue). E-F) Relative electron transfer rate of PSII (rETR) (E) and NPQ (F) during RLC measurement from dark-acclimated ambient-air (black solid lines) and high-CO_2_ *E. timida* (blue dashed lines) after 10 days in fluctuating-light starvation. Illumination for each light step during the RLCs was continued for 90 s prior to firing a saturating pulse to estimate rETR and NPQ. The light regime during the fluctuating light starvation was 12/12h day/night, and PPFD of the background illumination was 40 µmol m^-2^s^-1^, which was supplemented every 10 min with a 10 s high-light pulse during daylight hours. All data in (A,B) represent averages from 45 to 20 slug individuals (see “Materials and methods” for details on sampling). P700 redox kinetics in (C) represent averages from 9 biological replicates for both ambient-air and high-CO_2_ *E. timida*, and 6 and 9 in (D), respectively. P700^+^ transients were double normalized to their respective dark levels and to the P700^+^ peak measured immediately after the onset of the pulse. Fluorescence based data in (E,F) represent averages of 5 biological replicates for ambient-air and high-CO_2_ *E. timida*. All error bars and shaded areas around the curves show SE. See Figure 7 – source data 1 for original data.

Our results suggest that alternative electron acceptors of PSI, likely FLVs, are utilized by *E. timida* to protect kleptoplasts from formation of ROS during fluctuating-light starvation. The exact mechanism of PSI damage is not clear, but FLVs have been shown to protect PSI in green algae by donating excess electrons to oxygen without producing ROS (Shimakawa et al., 2019; Jokel et al., 2018). However, both slug groups did retain PSI activity for at least up to 6 days in starvation (Figure 7D). This shows that PSI was protected against fluctuating-light even in the high-CO_2_ *E. timida* that exhibited lowered, but not completely abolished, P700 oxidation capacity. Lower P700 oxidation capacity by FLVs in high-CO_2_ slugs could, however, cause an increase in the rate of Mehler’s reaction (Mehler, 1951;, Khorobrykh et al., 2020). Superoxide anion radical and hydrogen peroxide, the main ROS in Mehler’s reaction, are not likely to be involved in the primary reactions of PSII photoinhibition but are known to have deleterious effects on PSII repair (Tyystjärvi, 2013). We propose that this is behind the faster decrease in F_V_/F_M_ and rETR in the high-CO_2_ *E. timida* in fluctuating light.

## Discussion

We have performed the most detailed analysis of the differences in photosynthetic light reactions between a photosynthetic sea slug and its prey alga to date. Our results indicate that in the dark the PQ pool of the kleptoplasts inside the sea slug *E. timida* is not reduced to the same extent as in chloroplasts inside the green alga *Acetabularia* (Figure 2). We interpret this as a possible sign of a missing contribution of respiratory electron donors into the kleptoplasts inside *E. timida*. Either *E. timida* kleptoplasts are completely cut off from respiratory electron donors deriving from the slug’s mitochondria, or they are not delivered into the PQ pool due to inhibition of the NDH complex.

Fluorescence induction measurements also suggest that there are differences in the PQ pool redox state between kleptoplasts in *E. timida* and chloroplasts in *Acetabularia*. The considerable delay in reaching the maximum chlorophyll *a* fluorescence in *E. timida* during a high-light pulse (Figure 3A) can be indicative of a highly oxidized PQ pool that simply takes longer to fully reduce. Maintaining an oxidized PQ pool could be advantageous for chloroplast longevity, as it could help prevent electron pressure and ROS formation in PSII.

The strong build-up of proton motive force during transition from dark to light (Figure 4A) is most probably behind the fast induction and high levels of NPQ in *E. timida* (Figure 5G). Such alterations to photoprotective mechanisms provide an obvious benefit for long term maintenance of the kleptoplasts. However, the build-up also implies that protons do not diffuse out of the thylakoid lumen in *E. timida* kleptoplasts as efficiently as in *Acetabularia*. Further investigations into the function of ATP-synthase in *E. timida* could provide insights into the interchange of important molecules such as phosphate between the slug cells and kleptoplasts.

Our results show that both *E. timida* and *Acetabularia* utilize oxygen-dependent electron acceptors of PSI during dark-to-light transition (Figure 4B-C). Based on the current literature, the most likely candidates for these electron acceptors are FLVs (Shimakawa et al., 2019; Gerotto et al., 2016; Jokel et al., 2018). Oxidation of P700 by FLVs seems to be weaker in *E. timida* than in *Acetabularia* (Figure 4B, Figure 5 A-D), but FLVs do offer protection from light-induced damage in fluctuating light in *E. timida* (Figure 7). If the capacity to oxidize P700 by alternative electron acceptors is lowered in *E. timida* kleptoplasts, is this compensated for by an increased capacity of the main electron sink, i.e. the Calvin-Benson-Bassham cycle? If not, *E. timida* slugs would risk having a foreign organelle inside their own cells that readily produces ROS via one-electron reduction of oxygen. Interestingly, a major feature separating Sacoglossan slug species capable of long-term retention of kleptoplasts from those that are not, is their high capacity to downplay starvation induced ROS accumulation (de Vries et al., 2015). This could imply that long-term retention slug species such as *E. timida* do not need to concern themselves over the perfect functionality of the electron transfer reactions downstream of PSI. Further in-depth investigations into the carbon fixation reactions in photosynthetic sea slugs are needed to test this hypothesis. In addition to bringing closure to a biological conundrum that has remained unanswered for decades, solving how sea slugs are able to incorporate and maintain kleptoplasts in their own cells could provide useful insights into the ancient endosymbiotic events that led to the evolution of eukaryotic life.

## Materials and methods

### Organisms and culture conditions

Axenic stock cultures of the green alga *Acetabularia* (Düsseldorf Isolate 1, DI1; strain originally isolated by Diedrik Menzel) were grown in 5-10 l plastic tanks in sterile filtered f/2 culture medium made in 3.7 % artificial sea water (ASW; Sea Salt Classic, Tropic Marin). In order to slow down the stock culture growth, PPFD of growth lights (TL-D 58W/840 New Generation fluorescent tube; Philips, Amsterdam, The Netherlands) was <20 µmol m^-2^s^-1^. The culture medium for the stock cultures was changed at 8-10 week intervals. Other *Acetabularia* culture maintenance procedures, such as induction of gamete release, formation of zygotes and sterilization procedures were performed essentially as described earlier (Hunt and Mandoli, 1992; Cooper and Mandoli, 1999). The day-night cycle was 12/12 h and temperature was maintained at 23 °C at all times for all algae and slug cultures, unless mentioned otherwise. Algae used in the experiments were transferred to new tanks containing fresh f/2 media and grown under lights adjusted to PPFD 40 µmol m^-2^s^-1^ (TL-D 58W/840 New Generation fluorescent tube) for minimum of two weeks prior to any further treatments. No attempt was made to use only algae of certain age or size, and all populations were mixtures of cells in different developmental stages. PPFD was measured with a planar light sensor (LI-190R Quantum Sensor; LI-COR Biosciences; Lincoln, NE, USA) at the tank bottom level in all growth and treatment conditions. Irradiance spectra of all growth light sources used in the current study are shown in Figure 6 – figure supplement 1, measured with an absolutely calibrated STS-VIS spectrometer (Ocean Optics, Largo, FL, USA).

Sea slug *E. timida* individuals (50 individuals in total) were initially collected from the Mediterranean (Elba, Italy, 42.7782° N, 10.1927° E). The slug cultures were routinely maintained essentially as described by Schmitt et al. (2014). Briefly, *E. timida* were maintained at the same conditions as the cultures of *Acetabularia*, their prey alga, in aerated 5-10 l plastic tanks containing 3.7 % ASW. Fresh ASW was added to the tanks weekly to account for evaporation, and the slugs were placed in new tanks with fresh ASW at 3-5 week intervals. Differing amounts of *Acetabularia* were added to the slug tanks at irregular intervals, usually once every two weeks. When the adult slugs were transferred to new tanks, the old tanks with their contents were not discarded but supplemented with fresh ASW media and *Acetabularia* in order to allow unhatched slugs or slugs still in their veliger stage to develop into juvenile/adult slugs that are visible to the eye and could be pipetted out with a 10 ml plastic Pasteur pipette. The development from microscopic veligers to juvenile slugs usually took 2-3 weeks. Our method for cultivating slugs has enabled us to maintain a constant slug population consisting of 500-1000 slugs with relatively little labour and cost for years. It is, however, difficult to maintain the slug cultures axenic, and the slug tanks do foul, if not attended to. The contaminants in our laboratory cultures have not yet been identified but, based on optical inspection, seem to consist mainly of diatoms and ciliates. These organisms are likely derived from the Mediterranean and have been co-cultured with the slugs throughout the years. All slugs used in the experiments were always transferred into new tanks filled with fresh ASW and fed with abundant *Acetabularia* for 1-2 weeks prior to the experiments, unless mentioned otherwise. Slug individuals taken straight from the normal culture conditions were used for some of the measurements, without special considerations on the retention status of the chloroplasts inside the slugs, i.e. the slugs were not allowed to incorporate the chloroplasts overnight in the dark. The use of slugs without overnight settling time is indicated in the figures.

Acclimating *Acetabularia* to elevated CO_2_ levels was done in a closed culture cabinet (Algaetron AG230; Photon Systems Instruments, Drásov, Czech Republic) by raising the CO_2_ level from the ambient concentration (0.04 % of air volume) to 1 % CO_2_ inside the cabinet. Plastic 5 l tanks filled with *Acetabularia* were placed inside the cabinet and the tank lids were slightly opened to facilitate gas exchange. Fresh f/2 medium was added every second day to account for evaporation. Incident light provided by the growth cabinet white LEDs (see Figure 6 – figure supplement 1A for the spectrum) was adjusted to PPFD 40 µmol m^-2^s^-1^ and the day-night cycle was 12/12 h. Temperature was maintained at 23 °C. The algae were always acclimated for a minimum of three days to high CO_2_ prior to any measurements or feeding of the slugs with high-CO_2_ acclimated algae.

The red morphotype of *Acetabularia* was induced by growing the algae at 10 °C and continuous high white light (PPFD 600 µmol m^-2^s^-1^) for 31 days in a closed culture cabinet in ambient air (Algaetron AG230; Photon Systems Instruments), essentially as described by Costa et al. (2012). While this procedure was successful in producing the desired colour morphotype of *Acetabularia*, the yield was very low, and most of the *Acetabularia* cells bleached during the treatment. A few cells of red *Acetabularia* could be found in tanks where the cell concentration had been high enough to create a light attenuating algal mat. These red *Acetabularia* cells were then collected and used for measurements or fed to the slugs as indicated.

### Fast kinetics of Q_A_^-^ reoxidation, fluorescence induction (OJIP), P700 oxidation and ECS

Algae and slug samples were dark acclimated for 1-2 h prior to the fast kinetics measurements and all fast kinetics were measured at room temperature. PSII electron transfer was blocked in certain measurements, as indicated in the figures, with DCMU. For this, a 2 mM stock solution of DCMU in dimethylsulfoxide was prepared and diluted to a final concentration of 10 µM in either f/2 or ASW medium, depending on whether it was administered to the algae or the slugs, respectively. DCMU was only applied to samples that had been in the dark for 1 h and the dark acclimation in the presence of DCMU was continued for additional 20 min. When pertinent, DCMU containing medium was applied to cover the samples during the actual measurements too.

Anaerobic conditions were achieved by a combination of glucose oxidase (8 units/ml), glucose (6 mM) and catalase (800 units/ml) in f/2 or ASW medium. Our data shows that using the above reagents and concentrations in a sealed vial with stirring, nearly all oxygen was consumed from 250 ml of ASW media in a matter of minutes (Figure 2 – figure supplement 1A). Similar to the DCMU treatments, anaerobic conditions were initiated only after the samples had been in the dark for 1 h. In the case of *Acetabularia*, the samples were placed inside a sealed 50 ml centrifuge tube filled with f/2 medium that had been pre-treated with the glucose oxidase system for 10 min. The algae were then kept inside the sealed tube for 5 min in the dark, after which they were picked out and placed to the sample holder of the instrument in question. In order to maintain oxygen concentrations as low as possible, the samples were then covered with the anaerobic medium and left in the dark for additional 5 min, so that the oxygen mixed into the medium during sample placement would be depleted. Before imposing anaerobic conditions to slug individuals, they were swiftly decapitated with a razor blade, a procedure that has been shown not to significantly affect PSII activity in the photosynthetic sea slug *Elysia viridis* during a 2 h measurement period (Cruz et al., 2015). The euthanized slugs were then treated identically to the algae used in the anaerobic measurements.

Q_A_^-^ reoxidation kinetics after a strong single turnover (ST) flash (maximum PPFD 100 000 µmol m^-2^s^-1^, according to the manufacturer) were measured using an FL 200 fluorometer with a SuperHead optical unit (Photon Systems Instruments, Drásov, Czech Republic) utilizing the software and protocol provided by the manufacturer. The measurement protocol was optimized to be robust enough to allow its use in measurements from both *Acetabularia* and the slugs. The parameters used in the script were as follows: experiment duration - 120 s, Number of datapoints/decade - 8, First datapoint after ST flash - 150 µs, ST flash voltage – 100 %, ST flash duration – 30 µs, measuring beam (MB) voltage – 60 %. The wavelength for the ST flash and the MB was 625 nm. The option to enhance the ST flash intensity by complementing it with the MB light source was not used in the measurements. Number of datapoints/decade was changed to 2 for the measurements in the presence of DCMU.

The slugs tend to crawl around any typical cuboid 2 ml measuring cuvette if the cuvette is filled with ASW, which causes disturbances to the fluorescence signal. On the other hand, if ASW is removed from the cuvette, the slugs tend to stick to the bottom, placing them away from the light path of the instrument. For this reason, a compromise was made between ideal optics and slug immobilization by placing 3-5 slugs into a regular 1.5 ml microcentrifuge tube and then pipetting most of the ASW out of the tube, leaving just enough ASW to cover the slugs. Only in the measurements in anaerobic conditions were the tubes filled with oxygen depleted ASW. The tube was placed into the cuvette holder of the SuperHead optical unit so that the narrow bottom of the tube with the slugs was situated in the middle of the light path of the instrument and the tube was resting on its top appendices. The tube caps were left open for the measurements without any inhibitors and in the presence of DCMU, unlike the anaerobic measurements where the caps were closed. In the context of Q_A_^-^ reoxidation data from the slugs, one biological replicate refers to one measurement from 3-5 slugs inside the same tube in this study. Completely new slugs were used for each biological replicate. In order to facilitate comparison, Q_A_^-^ reoxidation from the algae was also measured using 1.5 ml microcentrifuge tubes, but due to the sessile nature of the algae there was no need to remove the f/2 media from the tubes. For each biological replicate representing the algae in the Q_A_^-^ reoxidation data sets, approximately 5-10 cells were placed inside each of the tubes.

The polyphasic fluorescence rise kinetics (OJIP curves) were measured with AquaPen-P AP 110-P fluorometer (Photon Systems Instruments) that has an inbuilt LED emitter providing 455 nm light for the measurements. The fluorometer was mounted on a stand and all measurements were done by placing a Petri dish with the sample on it on a matte black surface and positioning the sample directly under the probe head of the fluorometer. The intensity of the 2 s multiple turnover (MT) saturating pulse used for the measurements was optimized separately for measurements from single slug individuals (representing one biological replicate) and 1-2 cells/strands of algae (representing one biological replicate) placed under the probe of the instrument.

The final MT pulse intensity setting of the instrument was 70 % (100 % being equal to PPFD 3000 µmol m^-2^s^-1^ according to the manufacturer’s specifications) for the slugs and 50 % for *Acetabularia*. OJIP curves were measured from samples that had been covered in their respective treatment media, which presented a concern with regard to the anaerobic measurements, as the samples were not in a closed environment and oxygen diffusion into the sample could not be prevented. The data in Figure 2 – figure supplement 1B shows that diffusion is largely negated during the additional 5 min dark period even in an open setup. The conditions during these OJIP measurements will be referred to as anaerobic although some diffusion of oxygen to the samples occurred.

Fast kinetics of P700 oxidation during a 780 ms MT pulse were measured with Dual-PAM 100 (Heinz Walz GmbH, Effeltrich, Germany) equipped with the linear positioning system stand 3010-DUAL/B designed for plant leaves and DUAL-E measuring head that detects absorbance changes at 830 nm (using 870 nm as a reference wavelength). The absorbance changes are not caused entirely by P700 redox state, as the contribution of other components of the electron transfer chain, plastocyanin and ferredoxin, cannot be distinguished from the P700 signal at this wavelength region (Klughammer and Schreiber, 2016). P700 measurements were carried out essentially as described by (Shimakava et al., 2019), with slight modifications. We built a custom sample holder frame that can be sealed from the top and bottom with two microscope slide cover glasses by sliding the cover glasses into the. The frame of this sample holder was wide enough so that the soft stoppers of the Dual-PAM detector unit’s light guide could rest on it without disturbing the sample even when the top cover glass was not in place. A 3D-printable file for the sample holder is available at https://seafile.utu.fi/d/2bf6b91e85644daeb064/.

The 635 nm light provided by the LED array of Dual-PAM was used for the MT pulse (780 ms, PPFD 10000 µmol m^-2^s^-1^) in the P700^+^ measurements. Fluorescence was not measured during the MT pulse, as the MB used for fluorescence seemed to disturb the P700^+^ signal from the slugs. The drift of the signal made attempts to estimate the maximum oxidation level of P700 according to the standard protocol described by Schreiber and Klughammer (2008a) impossible with the slugs. Measurements from the algae would not have required any special considerations due to a stronger signal, but the algae were nevertheless measured identically to the slugs for the sake of comparability. The measuring light intensity used for detecting the P700 absorbance changes had to be adjusted individually for each sample. All P700^+^ kinetics were measured from individual slugs (i.e one slug represents one biological replicate), as pooling multiple slugs together for a single measurement did not noticeably enhance the signal.

Due to the delicate nature of the P700^+^ signal, all slugs used for these measurements had to be decapitated with a razor blade before the measurements. It is important to note that obtaining a single, meaningful fast kinetics curve of P700 oxidation requires sacrificing a lot of slug individuals. In this study a minimum of 10 individuals were used to construct each curve, because in approximately 30-50 % of the measurements the signal was simply too noisy and drifting to contain any meaningful information.

The P700^+^ measurements were carried out similarly to the OJIP measurements, with three main differences. First, the number of algae cells per measurement (representing one biological replicate) was higher, usually 5-10 cells/strands forming an almost solid green area between the light guides of Dual-PAM inside the sample holder. Secondly, the anaerobic measurements were carried out in a sealed system, achieved by closing the sample holder with both cover glasses after filling it with anaerobic medium. Measurements from all other treatments were carried out in open sample holders. The third difference was that for some of the experiments a second MT pulse was fired after a 10 s dark period following the first MT pulse. This procedure is referred to as “second pulse P700 redox kinetics protocol” in the main text.

Electrochromic shift (ECS, or P515) during a MT pulse (780 ms, 635 nm, PPFD 10000 µmol m^-2^s^-1^) was measured with P515 module of Dual-PAM 100 using the dual beam 550-515 transmittance difference signal (actual wavelengths used were 550 and 520 nm) (Schreiber and Klughammer, 2008b; Klughammer et al., 2013). ECS from *Acetabularia* was measured using the exact same setup as with the P700 measurements, but ECS from the slug *E. timida* could only be measured using the pinhole accessory of Dual-PAM 100. Shortly, a pinhole plug was placed on the optical rod of the P515 detector and 3-5 decapitated slug individuals (representing one biological replicate) were placed into the hole of the plug, covering the optical path. After placing a sample between the optical rods of the P515 module, the ECS signal was calibrated and the MB was turned off to decrease the actinic effect caused by the MB. MB was turned back on again right before measuring the ECS kinetics during a MT pulse. The intensity of the MB was adjusted for each sample separately.

The P700 oxidation and ECS data from *E. timida* and *Acetabularia* were slope corrected, when needed, using the baseline subtraction tool of Origin 2016 v.9.3 (OriginLab Corporation, Northampton, MA, USA) to account for signal drift. All biological replicates used to construct the fast kinetics data figures (Q_A_^-^ reoxidation, OJIP, P700 oxidation and ECS) were normalized individually as indicated in the main text figures, and the normalized data were averaged to facilitate comparison between the samples.

### Maximum quantum yield of PSII and rapid light response curves

Maximum quantum yield of PSII photochemistry (F_V_/F_M_) was routinely measured from slug individuals using PAM-2000 fluorometer (Heinz Walz GmbH) after minimum of 20 min darkness. The measurements were carried out by placing a dark acclimated slug on to the side of an empty Petri dish and then pipetting all ASW media out, leaving the slug relatively immobile for the time required for the measurement. The light guide of PAM-2000 was hand-held at a ∼45° angle respective to the slug, using the side and bottom of the Petri dish as support, and a saturating pulse was fired. PAM-2000 settings used for F_V_/F_M_ measurements from the slugs were as follows: MB intensity 10 (maximum), MB frequency 0.6 kHz, high MB frequency 20 kHz (automatically on during actinic light illumination), MT pulse intensity 10 (maximum, PPFD >10 000 µmol m^-2^s^-1^), MT pulse duration 0.8 s.

Measuring rapid light curves (RLCs) requires total immobilization of the slugs, a topic that has been thoroughly discussed by Cruz et al. (2012). Instead of using the anaesthetic immobilization technique described by Cruz et al. (2012), we tested yet another immobilization method to broaden the toolkit available for studying photosynthesis in Sacoglossan sea slugs. Alginate is a porous, biologically inert and transparent polymer that is widely used for fixing unicellular algae and cyanobacteria to create uniform and easy to handle biofilms or beads in e.g. biofuel research (Kosourov and Seibert, 2009; Antal et al., 2014). For the fixation of the slugs, an individual slug (representing one biological replicate) was placed on a Petri dish and a small drop of 1 % alginate (m/v in H_2_O) was pipetted on top of the slug, covering the slug entirely. Next, roughly the same volume of 0.5 mM CaCl_2_ was distributed evenly to the alginate drop to allow the Ca^2+^ ions to rapidly polymerize the alginate. The polymerization was allowed to continue for 10-30 s until the alginate had visibly solidified. All leftover CaCl_2_ was removed with a tissue, and the slug fixed inside the alginate drop was placed under the fixed light guide of PAM-2000, in direct contact and in a 90 ° angle, for the measurement. After the measurement was over, the alginate drop was covered with abundant 1M Na-EDTA to rapidly chelate the Ca^2+^ ions and depolymerize the alginate. Once the slug was visibly free of alginate, it was immediately transferred to fresh ASW for rinsing with a Pasteur pipette. The slugs usually recovered full movement, defined as climbing the walls of the container, in 10-20 min. The slugs were placed into a new tank for breeding purposes once motility had been restored. We also tested the effect of alginate fixation on F_V_/F_M_ during a 10 min time period, a typical length for RLC measurements, and no effect was noticeable (Figure 5 – figure supplement 1). All RLCs from algae were measured from 5-10 cells/measurement (representing one biological replicate), using otherwise the same setup as with the slugs, except that alginate fixation was not applied. The basic settings for RLC measurements were the same as with F_V_/F_M_ measurements, except for the MB intensity, which was adjusted to setting 5 with the algae to avoid oversaturation of the signal. Each light step lasted 90 s and the PPFDs that were used are shown in the figures. rETR was calculated as 0.42*Y(II)*PPFD, where 0.42 represents the fraction of incident photons absorbed by PSII, based on higher plant leaf assumptions, and Y(II) represents effective quantum yield of PSII photochemistry under illumination. NPQ was calculated as F_M_/F_M’_-1, where F_M’_ represent maximum chlorophyll fluorescence of illuminated samples. See Kalaji et al. (2014) for detailed descriptions of rETR and NPQ.

### Feeding experiments

In order to ensure that the slugs incorporate only specifically acclimated chloroplasts inside their own cells, the first feeding experiment was done with slug individuals that had been kept away from their food for four weeks in 5 l tanks filled with fresh ASW medium in their normal culture conditions. The coloration of the slugs was pale after the starvation period, indicating a decrease in the chloroplast content within the slugs. Altogether 107 starved slug individuals were selected for the feeding experiments and divided to two tanks filled with abundant *Acetabularia* in f/2 culture medium, one tank containing high-CO_2_ acclimated algae (54 slugs) and the other one algae grown in ambient air (53 slugs). The tanks with the slugs and algae in them were put to their respective growth conditions for 4 days to allow the slugs to incorporate new chloroplasts inside their cells. The tanks were not aerated during the feeding, but the tank lids were open for both feeding groups. The elevated CO_2_ level in the closed culture cabinet posed a problem, as it noticeably affected the slug behaviour by making them sessile in comparison to normal growth conditions, probably due to increased replacement of O_2_ by CO_2_. Because of this, the slugs selected for feeding on the high-CO_2_ acclimated algae were fed in cycles where the tanks were in the closed CO_2_ cabinet for most of the time during the daylight hours, but taken out every few hours and mixed with ambient air by stirring and kept in the ambient-air conditions for 1-2 hours before taking the tanks back to the high-CO_2_ cabinet. The tanks were always left inside the closed cabinets for the nights in order to inflict minimal changes to the acclimation state of the algae. After 4 days of feeding, 50 slug individuals of similar size and coloration were selected from both feeding groups and distributed into 2 new 5 l tanks/feeding group, filled with approximately 2 l of 3.7 % ASW medium and containing no algae. All slugs (25+25 ambient-air slugs, 25+25 high-CO_2_ slugs) were moved to ambient-air growth conditions and kept in the dark overnight in order to allow maximal incorporation of the chloroplasts before starting the starvation experiment in steady-light conditions.

For the second feeding experiment the four-week pre-starvation period of the slugs was discarded to see whether the differences in photosynthetic parameters between the two feeding groups could be inflicted just by allowing the slugs to replace their old kleptoplasts with the specific chloroplasts fed to them. We selected 100 slugs from normal growth conditions and divided them once again into two tanks containing f/2 culture medium and *Acetabularia* that had been acclimated to ambient air (50 slugs) or high CO_2_ (50 slugs). The slugs were allowed to feed for 6 days, but otherwise the feeding protocol was identical to the one used in the first feeding experiment. After the sixth day of feeding, 45 slug individuals of similar size and coloration were selected from both feeding groups and divided to 2 new 5 l tanks (20+25 ambient-air slugs, 20+25 1 % CO_2_ slugs) filled with approx. 2 l of ASW and kept overnight in the dark before starting the starvation experiment in fluctuating light.

A third feeding experiment was conducted in order to create the red morphotype of *E. timida* (González-Wangüemert et al., 2006; Costa et al., 2012). Slug individuals from normal growth conditions were selected and placed in Petri dishes filled with f/2 culture medium and abundant red *Acetabularia*. The slugs were allowed to eat the algae for 2 days in the normal culture conditions of the slugs prior to the measurements. Three Petri dishes were filled with just the red form *Acetabularia* in f/2 culture medium in exactly the same conditions, and these algae were used for measurements regarding the red morphotypes of *Acetabularia* and *E. timida*.

### Starvation experiments

Two different starvation experiments were carried out with ambient-air slugs and high-CO_2_ slugs. In the first one the slugs from the first feeding experiment were starved in steady-light conditions, where the only changes in the incident light were due to the day/night light cycle (12/12 h). Here, all four tanks (25+25 ambient-air slugs, 25+25 high-CO_2_ slugs) were placed under white LED lights (Växer PAR30 E27, 10 W; Ikea, Delft, The Netherlands; see Figure 6 – figure supplement 1B for the spectrum) adjusted to PPFD 40 µmol m^-2^s^-1^. Temperature was maintained at 23 °C and the tanks were not aerated during the starvation experiment apart from the passive gas flux that was facilitated by the open lids of the tanks. Fresh ASW medium (approximately 500 ml) was added to the tanks every second day and the slugs were placed into new tanks with fresh ASW 1-2 times a week throughout the entire starvation period of 46 days. The day following the overnight dark period that the slugs were subjected to after the feeding experiment was noted as day 0 in the starvation experiments. F_V_/F_M_ during starvation was measured from individual slugs (representing one biological replicate) as indicated in the main text figures.

Sampling caused losses to the slug populations on days 0, 5 and 15, when 10 slug individuals/group were selected for P700 oxidation kinetics measurements. Unfortunately, the P700^+^ signals after 15 days in starvation were too weak and noisy for any meaningful interpretations. Before day 25, starvation induced mortality of both groups was 0. After that the ambient-air slug population suffered losses on days 27 (1 slug), 35 (2 slugs), 45 (5 slugs), 46 (1 slug) altogether 9 slugs. For the high-CO_2_ slug population the losses were as follows: day 27 (1 slug), 29 (1 slug), 31 (1 slug), 45 (8 slugs) and 46 (1 slug), 12 slugs in total. Lengths of the slugs were estimated from images taken at set intervals essentially as described by Christa et al. (2018). Images were taken with a cropped sensor DSLR camera equipped with a macro lens (Canon EOS 7D MKII + Canon EF-S 60mm f/2.8 Macro lens; Canon Inc., Tokyo, Japan) and the body length of each slug individual was estimated using the open source image analysis software Fiji (Schindelin et al., 2012). The slugs from both feeding groups were pooled into one tank/group after day 25 in starvation.

The second starvation experiment was carried out using ambient-air slugs and high-CO_2_ slugs from the second feeding experiment. The four tanks from both feeding groups (20+25 ambient-air slugs, 20+25 high-CO_2_ slugs) were placed under a fluctuating light regime. The day/night cycle was maintained at 12/12 h, but during the daylight hours the background illumination (PPFD 40 µmol m^-2^s^-1^) was supplemented with a 10 s pulse of high light (PPFD 1500 µmol m^-2^s^-1^) every 10 minutes. Both the background illumination and the high-light pulses originated from a programmable Heliospectra L4A greenhouse lamp (model 001.010; Heliospectra, Göteborg, Sweden; see Figure 6 – figure supplement 1B for the irradiance spectra). All other conditions and procedures were identical to the ones used in the first starvation experiment. F_V_/F_M_ during starvation was measured as indicated in the main text figures. Sampling caused losses to the slug populations on days 0 and 6, when 10 slugs/group were selected for P700 redox kinetics measurements, and on day 10, when 5 slugs/group were selected for RLC measurements. No images were taken during the starvation experiment in fluctuating light. The slugs were pooled into one tank/feeding group after 25 days in starvation. Chlorophyll was extracted from the slugs with N,N-dimethylformamide and chlorophyll *a/b* was estimated spectrophotometrically according to Porra et al. (1989).

## Acknowledgments

This study was funded by Academy of Finland (grant 307335). V.H. was supported by Finnish Cultural Foundation, Väisälä Fund and University of Turku Graduate School (UTUGS). Sven. B. Gould and his research group are acknowledged for the invaluable help in establishing laboratory cultures of both *E. timida* and *Acetabularia*, and Taras Antal for help with fluorescence induction measurements.

## Figure supplements

**Figure 2- figure supplement 1.**
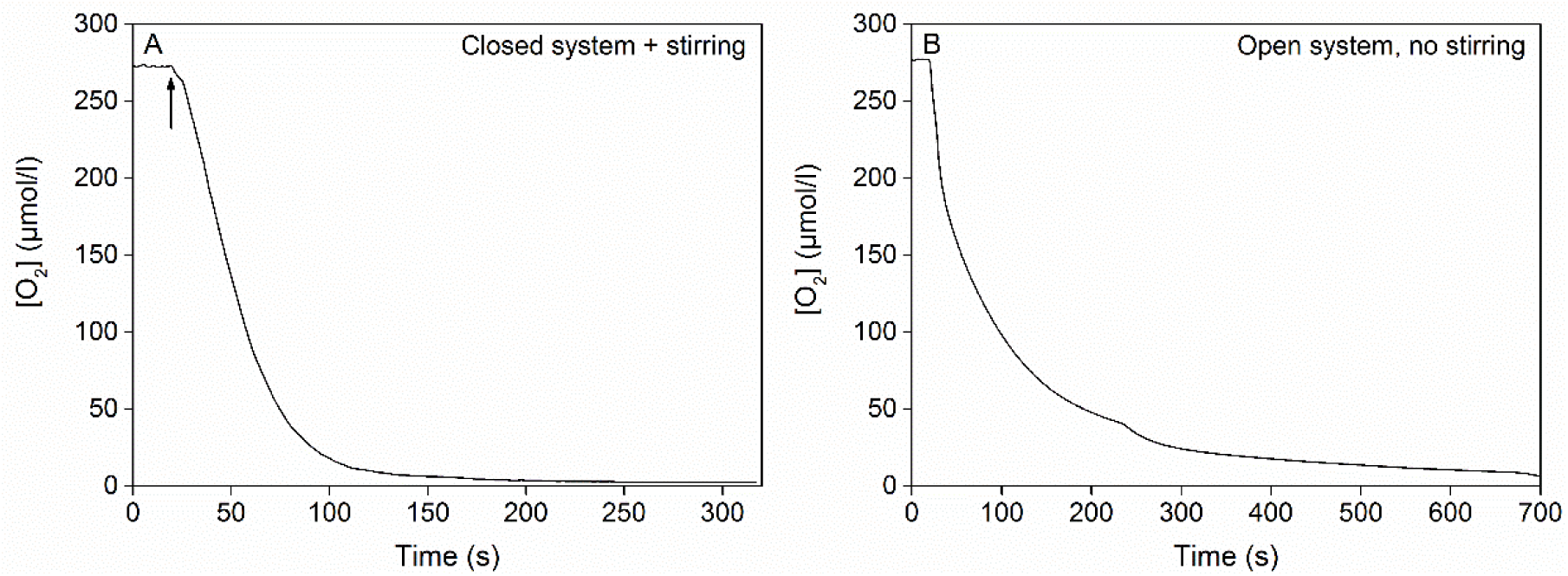
Oxygen consumption by the glucose oxidase system (8 units/ml glucose oxidase, 6 mM glucose and 800 units/ml catalase) in room temperature. A) Oxygen concentration in 250 ml of ASW medium inside a sealed bottle with minimal head space, stirred with a magnet, before and after the addition of glucose oxidase. The arrow indicates the point where the last component of the glucose oxidase system (glucose oxidase) was added into the mixture, after which the bottle was sealed. B) Oxygen concentration in 2 ml of ASW medium in an open cuvette without any stirring. The glucose oxidase system had been activated in a separate, sealed 5 ml vial 5 min prior to pipetting 2 ml of the activated mixture into an empty measuring cuvette. Oxygen concentration in (A) and (B) was measured with an optode-type oxygen meter FireStingO2 (PyroScience GmbH, Aachen, Germany) using optically isolated oxygen sensor spots according to manufacturer’s instructions.

**Figure 3 – figure supplement 1.**
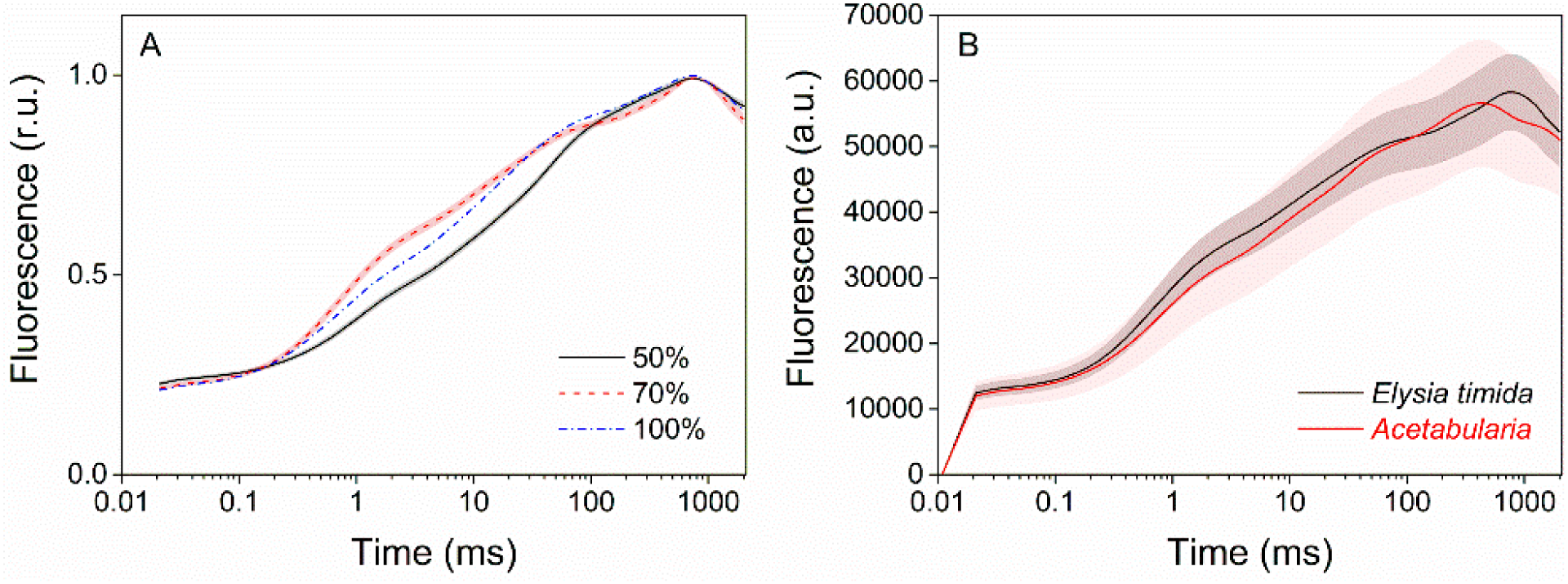
Technical considerations of the OJIP fluorescence induction measurements. A) Increasing the saturating pulse intensity from 50 % of the maximum (black solid line) to 70 % (red dashed line) or 100 % (PPFD 3000 µmol m^-2^s^-1^; blue dash-dot line) in *E. timida* measurements alters the O-J-I phases, but the intensity of the saturating pulse does not change the time required to reach maximum fluorescence. The data for the saturating pulse intensity 70 % are taken from Figure 3A. The curve for the 50 % saturating pulse represents an average from 12 biological replicates, and a representative curve is shown for the 100 % intensity measurement, because raising the saturating pulse intensity to >70 % often resulted in oversaturation of the fluorescence signal in *E. timida*. B) Original, unnormalized fluorescence traces from the data shown in Figure 3A, representing averages from 10 (*E. timida*) and 12 (*Acetabularia)* biological replicates. Shaded areas around the curves represent SE. All *E. timida* data are from individuals taken straight from the feeding tanks, without an overnight starvation period.

**Figure 5 – figure supplement 1.**
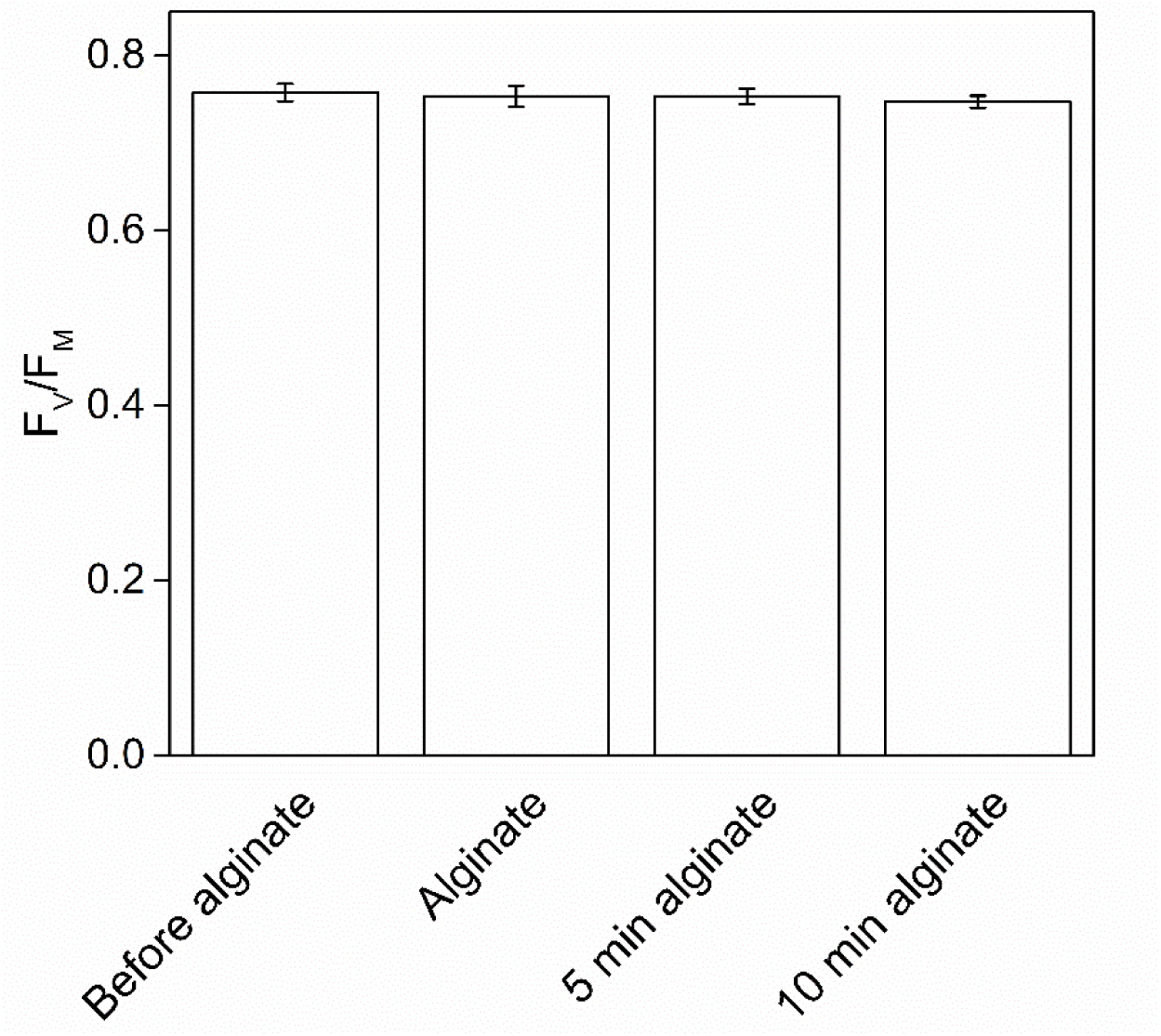
The effect of alginate fixation on the maximum quantum yield of PSII (F_V_/F_M_). Slug individuals were separately fixed in alginate and F_V_/F_M_ was monitored before the fixation, immediately after the fixation, and after 5 and 10 min in fixation. The slugs had been in the dark for 20 min before the first measurement and another 20 min dark period preceded the alginate fixation. The rest of the measurements were done at 5 min intervals, keeping the samples in the dark between the measurements. All data are averages from three biological replicates, and the error bars indicate SE.

**Figure 6 – figure supplement 1.**
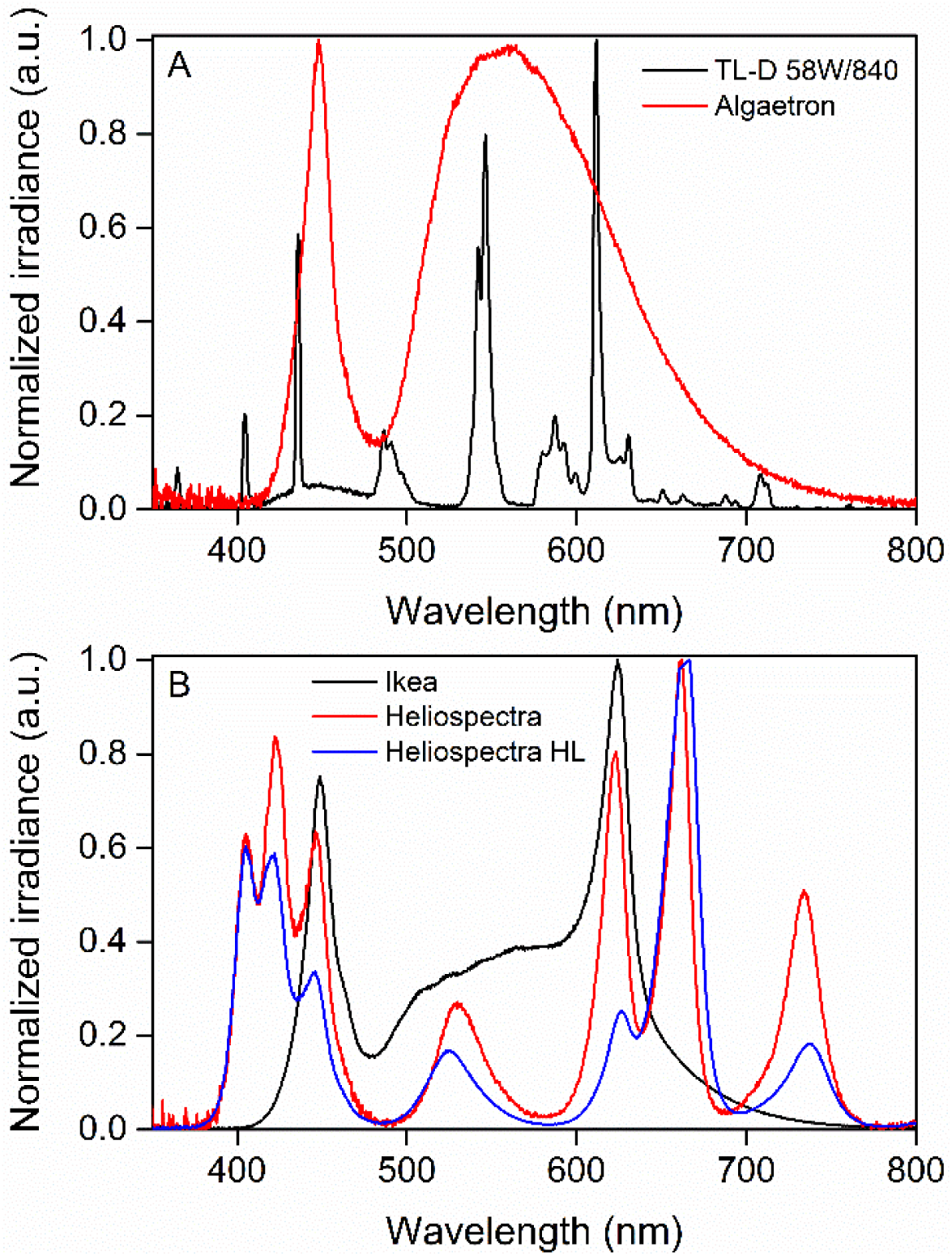
Normalized irradiance spectra from different light sources used in the study. A) Growth light spectra, TL-D 58W/840 New Generation fluorescent tube (black) and Algaetron AG230 LED array (red), used as illumination in regular growth conditions and during acclimation to high CO_2_, respectively. B) Light sources used in the starvation experiments: Ikea Växer PAR30 E27, 10 W (black) was used for the starvation experiment in steady light; Heliospectra L4A greenhouse lamp was used for the starvation experiment in fluctuating light, and the spectra are from moderate light conditions (PPFD 40 µmol m^-2^s^-1^; red) and during a high-light (HL) pulse (PPFD 1500 µmol m^-2^s^-1^; blue).

## Figure source data

Figure 2 – source data 1

Figure 3 – source data 1

Figure 4 – source data 1

Figure 5 – source data 1

Figure 6 – source data 1

Figure 7 – source data 1

## References

1. Alboresi A, Storti M, Cendron L, Morosinotto T (2019) Role and regulation of class-C flavodiiron proteins in photosynthetic organisms. Biochem. J. 476: 2487–2498. https://doi.org/10.1042/BCJ20180648

2. Allahverdiyeva Y, Isojärvi J, Zhang O, Aro EM (2015) Cyanobacterial Oxygenic Photosynthesis is Protected by Flavodiiron Proteins. Life 5: 716–743. https://doi.org/10.3390/life5010716

3. Antal TK, Matorin DN, Kukarskikh GP, Lambreva MD, Tyystjärvi E, Krendeleva TE, Tsygankov AA, Rubin AB (2014) Pathways of hydrogen photoproduction by immobilized *Chlamydomonas reinhardtii* cells deprived of sulfur. Int. J. Hydrog. Energy 39: 18194–1820. https://doi.org/10.1016/j.ijhydene.2014.08.135

4. Bulychev AA, Cherkashin AA, Muronets EM, Elanskaya IV (2018) Photoinduction of electron transport on the acceptor side of PSI in *Synechocystis* PCC 6803 mutant deficient in flavodiiron proteins Flv1 and Flv3. Biochim. Biophys. Acta 1859: 1086–1095. https://doi.org/10.1016/j.bbabio.2018.06.012

5. Cartaxana P, Morelli L, Jesus B, Calado G, Calado R, Cruz S (2019) The photon menace: kleptoplast protection in the photosynthetic sea slug *Elysia timida*. J. Exp. Biol. 222: jeb202580. https://doi.org/10.1242/jeb.202580

6. Cartaxana P, Trampe E, Kühl M, Cruz S (2017) Kleptoplast photosynthesis is nutritionally relevant in the sea slug *Elysia viridis*. Sci. Rep. 7: 7714. https://doi.org/10.1038/s41598-017-08002-0

7. Christa G, Pütz L, Sickinger C, Melo Clavijo J, Elaetz EMJ, Greve C, Serôdio J (2018) Photoprotective non-photochemical quenching does not prevent kleptoplasts from net photoinactivation. Front. Ecol. Evol. 6:121. https://doi.org/10.3389/fevo.2018.00121

8. Cooper JJ, Mandoli DF (1999) Physiological factors that aid differentiation of zygotes and early juveniles of *Acetabularia acetabulum* (Chlorophyta). J. Phycol. 35: 143–151. https://doi.org/10.1046/j.1529-8817.1999.3510143.x

9. Costa J, Giménez-Casalduero F, Melo RA, Jesus B (2012) Colour morphotypes of *Elysia timida* (Sacoglossa, Gastropoda) are determined by light acclimation in food algae. Aquat. Biol. 17: 81–89. https://doi.org/10.3354/ab00446

10. Cruz S, Cartaxana P, Newcomer R, Dionísio G, Calado R, Serôdio J, Pelletreau KN, Rumpho ME (2015) Photoprotection in sequestered plastids of sea slugs and respective algal sources. Sci. Rep. 5 : 7904. https://doi.org/10.1038/srep07904

11. Cruz S, Dionísio G, Rosa R, Calado R, Serôdio J (2012) Anesthetizing solar-powered sea slugs for photobiological studies. Biol. Bull. 223: 328–336. https://doi.org/10.1086/BBLv223n3p328

12. de Vries J, Habicht J, Woehle C, Huang C, Christa G, Wägele H, Nickelsen J, Martin WF, Gould SB (2013) Is ftsH the key to plastid longevity in sacoglossan slugs? Genome Biol. Evol. 5: 2540–8. https://doi.org/10.1093/gbe/evt205

13. de Vries J, Christa G, Gould SB (2014) Plastid survival in the cytosol of animal cells. Trends Plant Sci. 19: 347–350. https://doi.org/10.1016/j.tplants.2014.03.010

14. de Vries J, Woehle C, Christa G, Wägele H, Tielens AGM, Jahns P, Gould SB (2015) Comparison of sister species identifies factors underpinning plastid compatibility in green sea slugs. Proc. R. Soc. B. 282: 20142519. https://doi.org/10.1098/rspb.2014.2519

15. De Wijn R, van Gorkom HJ (2001) Kinetics of electron transfer from Q_A_ to Q_B_ in Photosystem II. Biochemistry 40: 11912–11922. https://doi.org/10.1021/bi010852r

16. Deák Z, Sass L, Kiss É, Vass I (2014) Characterization of wave phenomena in the relaxation of flash-induced chlorophyll fluorescence yield in cyanobacteria. Biochim. Biophys. Acta 1837: 1522–1532. https://doi.org/10.1016/j.bbabio.2014.01.003

17. Ermakova M, Huokko T, Richaud P, Bersanini L, Howe CJ, Lea-Smith DJ, Peltier G, Allahverdiyeva Y (2016) Distinguishing the roles of thylakoid respiratory terminal oxidases in the cyanobacterium *Synechocystis* sp. PCC 6803. Plant Physiol. 171: 1307–1319. https://doi.org/10.1104/pp.16.00479

18. Gerotto C, Alboresi A, Meneghesso A, Jokel M, Suorsa M, Aro EM, Morosinotto T (2016) Flavodiiron proteins as safety valve for electrons in *Physcomitrella patens*. Proc. Natl. Acad. Sci. USA 113: 12322–12327. https://doi.org/10.1073/pnas.1606685113

19. González-Wangüemert M, Francisca Giménez-Casalduero F, Perez-Ruzafa A (2006) Genetic differentiation of *Elysia timida* (Risso, 1818) populations in the Southwest Mediterranean and Mar Menor coastal lagoon. Biochem. Syst. Ecol. 34:514–527. https://doi.org/10.1016/j.bse.2005.12.009

20. Havurinne V, Mattila H, Antinluoma M, Tyystjärvi E (2018) Unresolved quenching mechanisms of chlorophyll fluorescence may invalidate multiple turnover saturating pulse analyses of photosynthetic electron transfer in microalgae. Physiol. Plant. 166: 365–379. https://doi.org/10.1111/ppl.12829

21. Hunt BE, Mandoli DF (1992) Axenic cultures of *Acetabularia* (Chlorophyta): A decontamination protocol with potential application to other algae. J. Phycol. 28: 407–414. https://doi.org/10.1111/j.0022-3646.1992.00407.x

22. Ilík P, Pavlovič A, Kouřil R, Alboresi A, Morosinotto T, Allahverdiyeva Y, Aro EM, Yamamoto H, Shikanai T (2017) Alternative electron transport mediated by flavodiiron proteins is operational in organisms from cyanobacteria up to gymnosperms. New Phytol. 214: 967–972. https://doi.org/10.1111/nph.14536

23. Järvi S, Suorsa M, Aro EM (2015) Photosystem II repair in plant chloroplasts - Regulation, assisting proteins and shared components with photosystem II biogenesis. Biochim. Biophys. Acta 1847: 900–9. https://doi.org/10.1016/j.bbabio.2015.01.006

24. Jokel M, Johnson X, Peltier G, Aro EM, Allahverdiyeva Y (2018) Hunting the main player enabling *Chlamydomonas reinhardtii* growth under fluctuating light. Plant J. 94: 822–835. https://doi.org/10.1111/tpj.13897

25. Kalaji HM, Schansker G, Ladle RJ, Goltsev V, Bosa K, Allakhverdiev SI, Brestic M, Bussotti F, Calatayud A, Dąbrowski P, Elsheery NI, Ferroni L, Guidi L, Hogewoning SW, Jajoo A, Misra AN, Nebauer SG, Pancaldi S, Penella C, Poli DB, Pollastrini M, Romanowska-Duda ZB, Rutkowska B, Serôdio J, Suresh K, Szulc W, Tambussi E, Yanniccari M, Zivcak M (2014) Frequently asked questions about *in vivo* chlorophyll fluorescence: practical issues. Photosynth. Res. 122: 121–158. https://doi.org/10.1007/s11120-014-0024-6

26. Khorobrykh S, Havurinne V, Mattila H, Tyystjärvi E (2020) Oxygen and ROS in photosynthesis. Plants 9: 91. https://doi.org/10.3390/plants9010091

27. Klughammer C, Schreiber U (2016) Deconvolution of ferredoxin, plastocyanin, and P700 transmittance changes in intact leaves with a new type of kinetic LED array spectrophotometer. Photosynth. Res. 128: 195–214. https://doi.org/10.1007/s11120-016-0219-0

28. Klughammer C, Siebke K, Schreiber U (2013) Continuous ECS-indicated recording of the protonmotive charge flux in leaves. Photosynth Res. 117: 471–487. https://doi.org/10.1007/s11120-013-9884-4

29. Kodru S, Malavath T, Devadasu E, Nellaepalli S, Stirbet A, Subramanyam R, Govindjee (2015) The slow S to M rise of chlorophyll *a* fluorescence reflects transition from state 2 to state 1 in the green alga *Chlamydomonas reinhardtii*. Photosynth. Res. 125:219–31. https://doi.org/10.1007/s11120-015-0084-2

30. Kosourov SN, Seibert M (2009) Hydrogen photoproduction by nutrient-deprived *Chlamydomonas reinhardtii* cells immobilized within thin alginate films under aerobic and anaerobic conditions. Biotechnol. Bioeng. 102: 50–8. https://doi.org/10.1002/bit.22050

31. Kramer DM, Di Marco G, Loreto F (1995) Contribution of plastoquinone quenching to saturation pulse-induced rise of chlorophyll fluorescence in leaves In: Mathis P (Ed.) Photosynthesis: From Light to Biosphere, Vol. 1, pp 147–150. Kluwer Academic Publishers, Dordrecht, The Netherlands.

32. Krishna PS, Morello G, Mamedov F (2019) Characterization of the transient fluorescence wave phenomenon that occurs during H_2_ production in *Chlamydomonas reinhardtii*. J. Exp. Bot. 70: 6321–6336. https://doi.org/10.1093/jxb/erz380

33. Magyar M, Sipka G, Kovács L, Ughy B, Zhu Q, Han G, Špunda V, Lambrev PH, Shen J, Garab G (2018) Rate-limiting steps in the dark-to-light transition of Photosystem II - revealed by chlorophyll-a fluorescence induction. Sci. Rep. 8: 2755. https://doi.org/10.1038/s41598-018-21195-2

34. Mehler AH (1951) Studies on reactivity of illuminated chloroplasts. Mechanism of the reduction of oxygen and other Hill reagents. Arch. Biochem. Biophys. 33: 65–77. https://doi.org/10.1016/0003-9861(51)90082-3

35. Müller P, Li XP, Niyogi KK (2001) Non-Photochemical Quenching. A Response to Excess Light Energy. Plant Physiol. 125: 1558–1566. https://doi.org/10.1104/pp.125.4.1558

36. Oja V, Eichelmann H, Laisk A (2011) Oxygen evolution from single- and multiple-turnover light pulses: temporal kinetics of electron transport through PSII in sunflower leaves. Photosynth. Res. 110: 99–109. https://doi.org/10.1007/s11120-011-9702-9

37. Osmond B, Chow WS, Wyber R, Zavafer A, Keller B, Pogson BJ, Robinson SA (2017) Relative functional and optical absorption cross-sections of PSII and other photosynthetic parameters monitored in situ, at a distance with a time resolution of a few seconds, using a prototype light induced fluorescence transient (LIFT) device. Funct. Plant. Biol. 44: 985–1006. https://doi.org/10.1071/FP17024

38. Papageorgiou GC, Govindjee (2014) The non-photochemical quenching of the electronically excited state of chlorophyll *a* in plants: Definitions, timelines, viewpoints, open questions. In: Demmig-Adams B, Garab G, Adams W III, Govindgee (Eds.) Non-photochemical quenching and energy dissipation in plants, algae and cyanobacteria. Advances in Photosynthesis and Respiration, Vol. 40, pp 1–44. Springer, Dordrecht, The Netherlands.

39. Porra RJ, Thompson WA, Kriedemann PE (1989) Determination of accurate extinction coefficients and simultaneous equations for assaying chlorophyll *a* and *b* with four different solvents: verification of the concentration of chlorophyll by atomic absorption spectroscopy. Biochim. Biophys. Acta 975: 384–394. https://doi.org/10.1016/S0005-2728(89)80347-0

40. Rauch C, Jahns P, Tielens AGM, Gould SB, Martin WF (2017) On being the right size as an animal with plastids. Front. Plant Sci. 8: 1402. https://doi.org/10.3389/fpls.2017.01402

41. Rumpho ME, Pelletreau KN, Moustafa A, Bhattacharya D (2011) The making of a photosynthetic animal. J. Exp. Biol. 214: 303–311. https://doi.org/10.1242/jeb.046540

42. Santana-Sanchez A, Solymosi D, Mustila H, Bersanini L, Aro EM, Allahverdiyeva Y (2019) Flavodiiron proteins 1–to-4 function in versatile combinations in O_2_ photoreduction in cyanobacteria. eLife 8: e45766. https://doi.org/10.7554/eLife.45766

43. Schansker G, Tóth SZ, Holzwarth AR, Garab G (2014) Chlorophyll *a* fluorescence: beyond the limits of the Q_A_ model. Photosynth Res. 120: 43–58. https://doi.org/10.1007/s11120-013-9806-5

44. Schindelin J, Arganda-Carreras I, Frise E, Kaynig V, Longair M, Pietzsch T, Preibisch S, Rueden C, Saalfeld S, Schmid B, Tinevez JY, White DJ, Hartenstein V, Eliceiri K, Tomancak P, Cardona A (2012) Fiji: an open-source platform for biological-image analysis. Nat. Methods 9: 676–682. https://doi.org/10.1038/nmeth.2019

45. Schmitt V, Händeler K, Gunkel S, Escande ML, Menzel D, Gould SB, Martin WF, Wägele H (2014) Chloroplast incorporation and long-term photosynthetic performance through the life cycle in laboratory cultures of *Elysia timida* (Sacoglossa, Heterobranchia). Front. Zool. 11: 5. https://doi.org/10.1186/1742-9994-11-5

46. Schreiber U, Klughammer C (2008a) Saturation pulse method for assessment of energy conversion in PSI. PAM Appl. Notes 1: 11–14.

47. Schreiber U, Klughammer C (2008b) New accessory for the DUAL-PAM-100: The P515/535 module and examples of its application. PAM Appl. Notes 1: 1–10.

48. Schreiber U, Klughammer C, Schansker G (2019) Rapidly reversible chlorophyll fluorescence quenching induced by pulses of supersaturating light *in vivo*. Photosynth. Res. 142: 35–50. https://doi.org/10.1007/s11120-019-00644-7

49. Shimakawa G, Murakami A, Niwa K, Matsuda Y, Wada A, Miyake C (2019) Comparative analysis of strategies to prepare electron sinks in aquatic photoautotrophs. Photosynth. Res. 139: 401–411. https://doi.org/10.1007/s11120-018-0522-z

50. Stirbet A, Govindjee (2012) Chlorophyll a fluorescence induction: a personal perspective of the thermal phase, the J-I-P rise. Photosynth. Res. 113: 15–61. https://doi.org/10.1007/s11120-012-9754-5

51. Strasser RJ, Srivastava A, Govindjee (1995) Polyphasic chlorophyll *a* fluorescence transient in plants and cyanobacteria. Photochem. Photobiol. 61: 32–42. https://doi.org/10.1111/j.1751-1097.1995.tb09240.x

52. Strasser RJ, Tsimilli-Michael M, Srivastava A (2004) Analysis of the chlorophyll *a* fluorescence transient. In: Papageorgiou GC, Govindjee (Eds.) Chlorophyll a Fluorescence: A Signature of Photosynthesis. Advances in Photosynthesis and Respiration, Vol. 19, pp 321–362. Springer, Dordrecht, The Netherlands.

53. Suggett DJ, Oxborough K, Baker NR, Macintyre HL, Kana TM, Geider RJ (2003) Fast repetition rate and pulse amplitude modulation chlorophyll *a* fluorescence measurements for assessment of photosynthetic electron transport in marine phytoplankton. Eur. J. Phycol. 38: 371–384. https://doi.org/10.1080/09670260310001612655

54. Tikkanen M, Grebe S (2018) Switching off photoprotection of photosystem I - a novel tool for gradual PSI photoinhibition. Physiol. Plant. 162: 156–161. https://doi.org/10.1111/ppl.12618

55. Tyystjärvi E (2013) Photoinhibition of Photosystem II. Int. Rev. Cell Mol. Biol. 300: 243–303. https://doi.org/10.1016/B978-0-12-405210-9.00007-2

56. Vredenberg W (2015) A simple routine for quantitative analysis of light and dark kinetics of photochemical and non-photochemical quenching of chlorophyll fluorescence in intact leaves. Photosynth. Res. 124:87–106. https://doi.org/10.1007/s11120-015-0097-x

57. Wang QJ, Singh A, Li H, Nedbal L, Sherman LA, Govindjee, Whitmarsh J (2012) Net light-induced oxygen evolution in photosystem I deletion mutants of the cyanobacterium *Synechocystis* sp. PCC 6803. Biochim. Biophys. Acta 1817: 792–801. https://doi.org/10.1016/j.bbabio.2012.01.004

58. Yaakoubd B, Andersen R, Desjardins Y, Samson G (2002) Contributions of the free oxidized and Q_B_-bound plastoquinone molecules to the thermal phase of chlorophyll-*a* fluorescence. Photosynth. Res. 74: 251. https://doi.org/10.1023/A:1021291321066

59. Zhang P, Eisenhut M, Brandt AM, Carmel D, Silén HM, Vass I, Allahverdiyeva Y, Salminen TA, Aro EM (2012) Operon *flv4-flv2* provides cyanobacterial photosystem II with flexibility of electron transfer. Plant Cell 24: 1952–71. https://doi.org/10.1105/tpc.111.094417

